# Development of a 24-Channel 3T Phased-Array Coil for fMRI in Awake Monkeys - Mitigating Spatiotemporal Artifacts in Ferumoxytol-Weighted Functional Connectivity Estimation

**DOI:** 10.1101/2025.04.15.648822

**Authors:** Joonas A. Autio, Atsushi Yoshida, Yoshihiko Kawabata, Masahiro Ohno, Kantaro Nishigori, Takayuki Ose, Stephen Smith, David C. Van Essen, Matthew F. Glasser, Takuya Hayashi

## Abstract

Functional magnetic resonance imaging (fMRI) of awake macaque monkeys holds promise for advancing our understanding of primate brain organization, including humans. However, estimating functional connectivity in awake animals is challenging due to the limited duration of imaging sessions and the relatively low sensitivity to neural activity. To overcome these challenges, we developed a 24-channel 3T receive radiofrequency (RF) coil optimized for parallel imaging of awake macaques. This enabled the acquisition of multiband and GRAPPA-accelerated ferumoxytol-weighted resting-state fMRI. The Human Connectome Project-style data processing pipelines were adapted to address the unique preprocessing demands of cerebral blood volume-weighted (CBVw) imaging, including motion correction, functional-to-structural image co-registration, and training a multi-run independent component analysis-based X-noiseifier (ICA-FIX) classifier for removal of structured artifacts. Our CBVw fMRI approach resulted in an elevated contrast-to-noise ratio compared to blood oxygenation level dependent (BOLD) imaging in anesthetized macaques. However, structured imaging artifacts still contributed more variance to the functional timeseries than neural activity. By applying the ICA-FIX classifier, we achieved highly reproducible parcellated functional connectivity at the single-subject level. At the group-level, we identified dense functional networks with spatial features homologous to those observed in humans. The developed RF receive coil, image acquisition protocols, and data analysis pipelines are publicly available, providing the broader scientific community with tools to leverage these advances for further research.

## 1. Introduction

The macaque monkey serves as a pivotal animal model in elucidating the intricate neuroanatomical and functional arrangements of the primate brain. Functional magnetic resonance imaging (fMRI) has become a fundamental tool in the investigation of default functional organization within both human and non-human primates (NHPs) (Hutchison et al., 2011; Hutchison & Everling, 2012; Smith et al., 2013; Vincent et al., 2007). Notably, comparison between resting-state and task-induced functional connectivity (FC) patterns in humans has also revealed a high degree of similarity, suggesting an inherent relationship between task-induced neural networks and resting-state brain activity (Glasser et al., 2018; Nickerson, 2018; Smith et al., 2009). Therefore, resting-state fMRI has the potential to bypass the major challenge of task-design across species, enabling evolutionary comparison of default functional organization among primates (Mantini et al., 2011; Xu et al., 2020; Yokoyama et al., 2021).

Despite the promise of interspecies comparisons, challenges persist due to the relatively limited sensitivity and specificity of fMRI, compounded by factors such as hardware constraints, anesthesia effects, and a lack of standardized data preprocessing pipelines, leading to reduced reproducibility in NHP resting-state fMRI studies (Autio et al., 2021; Botvinik-Nezer et al., 2020; Milham et al., 2018). To bridge this gap, recent advances have adapted human neuroimaging methodologies to NHPs, including the development of a specialized 3T multi-channel radiofrequency (RF) receive coils, facilitating the translation of imaging and data analysis strategies from the Human Connectome Project (HCP) to anesthetized NHPs (Autio et al., 2020; Donahue et al., 2018; Glasser et al., 2013; Hayashi et al., 2021; Ose et al., 2022; Smith et al., 2013). This neuroanatomically informed approach has a potential to significantly enhance our ability to map the structural and functional organization in non-human primates (NHPs) (Yokoyama et al., 2021). However, the influence of anesthesia on functional inter-species comparisons remains a concern (Paasonen et al., 2018; Vanduffel et al., 2001), prompting efforts to extend these NHP-HCP methodologies to awake macaque monkeys.

Imaging awake macaques introduces distinct challenges compared to studies with anesthetized subjects (Autio et al., 2021; Vanduffel et al., 2001). Notably, the necessity of surgically mounting a head-post for RF coil attachment alongside an animal chair must be balanced with practical constraints related to scan duration and potential discomfort. These constraints highlight the need for fast, accelerated imaging achievable through multiband (MB) imaging (Moeller et al., 2010), to acquire a large number of timepoints necessary for resting-state fMRI. Additionally, scan duration can also be eased by the application of an ultrasmall superparamagnetic iron oxide (USPIO) contrast agent, ferumoxytol, to boost the contrast-to-noise ratio (CNR) (Leite et al., 2002; Mandeville et al., 1999, 2004; Vanduffel et al., 2001), with optimal brain contrast achieved at the shortest feasible echo-time (TE) (Mandeville et al., 1999, 2004). The use of in-plane acceleration, such as generalized auto-calibrating partially parallel acquisitions (GRAPPA) (Griswold et al., 2002), further facilitates the reduction of TE and repetition time (TR). The combination of in- and out-of-plane accelerations set stringent demands on the parallel imaging performance of the RF coil (Autio et al., 2020; Gilbert et al., 2016).

The subsequent data analysis challenges of accelerated ferumoxytol-weighted fMRI also remain poorly understood. For instance, structured spatiotemporal artifacts impose a challenge which can significantly impact FC estimates (Power et al., 2012; Van Dijk et al., 2012). While the efficacy of independent component analysis-based X-noiseifier (ICA-FIX) artifact removal is well established in human BOLD fMRI studies (Glasser et al., 2018; Griffanti et al., 2014; Salimi-Khorshidi et al., 2014), its efficacy in CBVw fMRI experiments remains unexplored. For instance, ferumoxytol may partially wash-out during a functional imaging session, thereby introducing spurious global correlations across the brain. Moreover, it is well established that injection of contrast agents causes strong static magnetic field (B_0_) orientation bias (Autio et al., 2024; Ogawa et al., 1993). Yet, the impact of ferumoxytol to FC metrics remains largely unexplored.

In this study, we introduce a novel 24-channel 3T receive RF coil tailored for parallel imaging of awake macaque monkeys. Capitalizing on the parallel imaging capabilities of the phased-array RF receive coil, we acquire accelerated ferumoxytol-weighted resting-state fMRI in awake macaque monkeys. We explore the effect of ICA-FIX artifact cleanup to CBVw functional connectivity estimation and reproducibility of the within-and between macaques. We apply this approach to distinguish individual macaques by their unique neural fingerprints.

## 2. Methods

Experiments were performed using a 3T MRI scanner (MAGNETOM Prisma, Siemens, Erlangen, Germany) equipped with 80 mT/m gradients (XR 80/200 gradient system with slew rate 200 T/m/s) and a 2-channel B_1_ transmit array (TimTX TrueForm). The animal experiments were conducted in accordance with the institutional guidelines for animal experiments, and animals were maintained and handled in accordance with the institutional guidelines for animal experiments, Basic Policies for the Conduct of Animals Experiments in Research Institution (MEXT, Japan) and the Guide for the Care and Use of Laboratory Animals of the Institute of Laboratory Animal Resources (ILAR; Washington, DC, USA). All animal procedures were approved by the Animal Care and Use Committee of the Kobe Institute of RIKEN (MA2008-03-19, A2021-05-6).

### 2.1. Development of a Phased-Array 3T Receive-Coil for Awake, Headpost Implanted Macaque Monkeys

The 24-channel coil for awake macaque imaging presented here builds upon our previously published design (Autio et al., 2020). However, as noted in the introduction, imaging awake monkeys requires an implanted head-post, a secure attachment mechanism, and an MRI compatible MRI chair, features that were explicitly incompatible with our previous coil design.

Addressing this limitation was a non-trivial challenge from a coil design perspective. The headpost attachment necessitates a larger opening in the phased-array coil, which can potentially compromise SNR, channel decoupling, and parallel imaging performance. To mitigate these issues, we developed a modular, attachable coil design that accommodates the headpost while modified coil element reconfiguration maintains high SNR and parallel imaging acceleration capabilities. The attachable configuration enables both stability and flexibility in experimental setups while largely preserving the benefits of a dense phased-array coil for accelerated imaging.

#### 2.1.1 Coil Design, Construction, and Evaluation

The coil frame geometry was designed using a 3D digital design software (Rhinoceros5, McNeel, Seattle, USA) to closely fit over the headpost-implanted (*Macaca mulatta*) monkeys (Fig. 1A). The helmet-shaped inner coil frame was constructed from machine-carved polyphenylene sulfide (PPS) plates. The thin PPS plates (2 mm) minimized the distance between coil elements while providing good workability and resistance to the heat of soldering.

**Figure 1.**
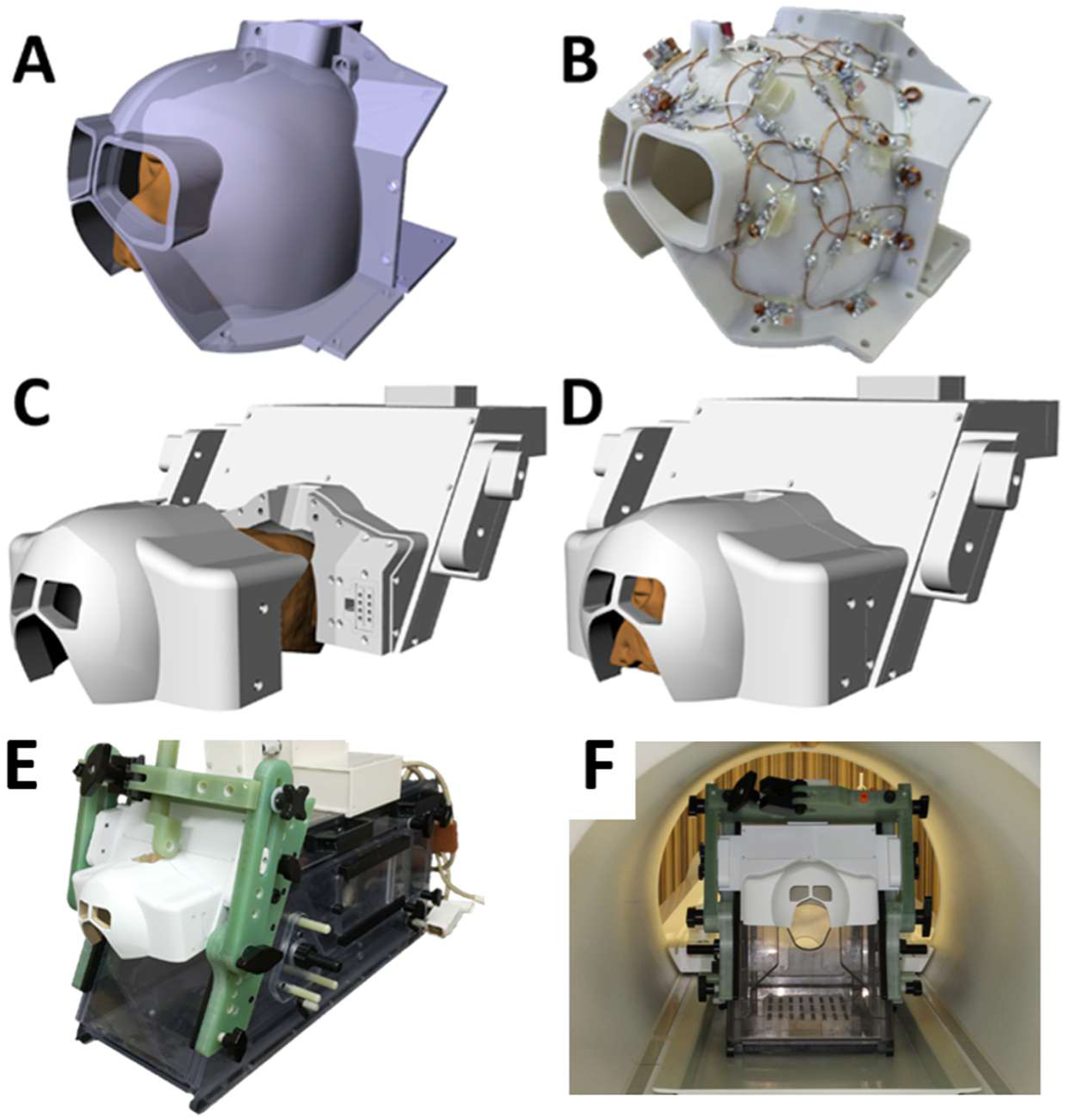
The design and development of a 24-channel receive coil for awake headpost implanted macaque MRI. **(A)** Design of coil geometry and **(B)** coil with 24-element arrangements. **(C, D)** The coil elements were separated into anterior and posterior halves to easily equip the coil on the monkey. **(E)** Coil attached to a customized MRI compatible primate chair (Rogue Research, Mtl, CA, Canada). (**F**) The primate chair and coil positioned on a customized plastic plate within the bore of the scanner. Note that the chair is placed on the plastic plate rather than on the scanner’s built-in bed.

Twenty-four coil elements were arranged over external surface, emulating a soccer ball coil design (G. c. Wiggins et al., 2006). To reduce potential interaction with RF transmit field (B_1_+), the elements were wired using thin coaxial copper cables (cable diameter 0.7 mm) and capacitors were arranged vertically against the coil surface (Autio et al., 2020; Wiggins et al., 2009). The coil elements covered the entire monkey’s head, excluding the mouth region, and were arranged with critical overlap to minimize coupling between neighboring elements (Supp. Fig. 1). This design improved decoupling between coil elements, which is essential for optimizing parallel encoding performance of multiband and GRAPPA accelerated MRI sequences. The elements were separated into anterior and posterior halves to more easily place the coil on the monkey head (Fig. 1C, D). The coil elements between two halves were overlapped by bending a part of elements outward and facing each other (Fig. 1B). The two elements arranged over the eyes were relatively large in diameter to allow video recording of eyes and eyelids for monitoring subject behavior (e.g., alertness vs drowsiness).

The outer cover (e.g. the frame of the coil; Fig. 1D) was designed for easy attachment to an MRI-compatible monkey chair (HypexInnovations, Rogue research, Inc., Canada) (Fig. 1E) and constructed using acrylic-based resin and a 3D printer (M200, Zortrax, Olsztyn, Poland). The chair was positioned on a custom-made plastic plate/holder, rather than being placed directly on the MRI bed (Fig. 1F). Overall, the resulting central location of the macaque brain (e.g. thalamus) was positioned approximately 10 cm above the B_0_ isocenter.

Circuit design followed standard design consisting of diode detuning trap, cable trap (to achieve common suppression for each element) and bias T connected to low input-impedance preamplifiers (Siemens Healthcare, Erlangen, Germany) (Wiggins et al, 2006, Autio et al., 2020).

The coil preamplifier (MPM series, Microwave Technology Inc.,CA, USA) was positioned above the monkey chair (Fig. 1E), with a cable length of 0.85 m from the coil to the preamplifier. Ideally, the preamplifier would be positioned adjacent to the coil to maximize SNR. However, such a configuration would significantly increase the coil size, making it impractical for experimental use. Pilot studies demonstrated that increasing the distance between the coil and preamplifier from directly adjacent to 1.5 m results in a 20% SNR reduction. To balance signal preservation with usability, we optimized the cable length to 0.85 m, corresponding to half of the RF wavelength specific to the cable material. This adjustment ensured impedance matching and phase consistency between the coil and preamplifier, minimizing signal loss.

Decoupling was achieved through a combination of geometrical and preamplifier decoupling. The latter was implemented by maintaining a very low impedance level in both the coil and at the preamplifier inputs. To ensure consistent impedance and phase alignment, the cable length was set to half the cable-specific RF wavelength, optimizing decoupling efficiency and preserving the performance of the 24-channel receive coil.

The coil elements were assessed for the ratio of loaded to unloaded quality factor Q, nearest-neighbor coupling, and active detuning, as well as gradient off-line noise correlation. The Q-ratio was assessed using a Vector Network Analyzer (E5061B, KeySight Technologies, CA, US) and a tightly fitted phantom (solution NiCl + NaCl) used for loading. The B_1_ transmit (B_1_+) field-map was estimated using a slice-selective RF-prepared sequence with a turbo fast low-angle-shot (FLASH) read-out (Chung et al., 2010). The B_1_ receive (B_1_-) field-map was estimated using a FLASH sequence, based on the signal intensity ratio between phased-array and body receive coils. Additionally, geometry-dependent noise amplification due to parallel imaging (g-factor) was assessed using gradient-echo imaging and GRAPPA and a phantom shaped to match the average macaque brain (Autio et al., 2020; Griswold et al., 2002).

#### 2.1.2. Headpost Surgery

Macaque monkeys (Macaca mulatta, N=3, weight=5.1 ± 0.8 kg, age=6.3 ± 1.0 y.o.) were trained to sit in a sphinx position in an MRI compatible primate chair (Rogue Research, Mtl, CA, Canada). Upon successful completion of the training, the animals were anesthetized and T1w images were acquired (see pilot study). The images were used to generate a bone contour of the subject’s skull surface (Rogue Research, Mtl, CA, Canada). Then, a custom-made headpost was produced to closely match the contour of each subject’s skull surface using PEEK (Poly Ether Ether Ketone, Rogue Research, Inc. Canada).

The headpost was surgically implanted in a sterile environment. The animals were initially anesthetized with ketamine (7.5 mg/ml/kg, i.m.) and xylazine (1.5 mg/ml/kg, i.m.), intubated and deep anesthesia was maintained using isoflurane (≥1.5%, Apollo anesthesia workstation, Dragerwerk AG & Co., Germany). Prior to surgery, the animals were administered antibiotics (Victus, DS Pharma Animal Health Co., Ltd, Japan; 5 mg/ml/kg, i.m.) and an analgesic (Sosegon, MaruishiPharmaceutical Co., Ltd, Japan; 0.2 ml, i.m.). The subjects’ heads were shaved and disinfected with isodine (10%, Meiji Seika Pharma Co., Ltd, Japan). During the surgery, body temperature was controlled with a heating pad (KN-474-S, Natsume Seisakusho, Co. Ltd, Tokyo, Japan) while physiological parameters (respiration rate, heart rate, oxygen saturation, body temperature, and expired CO_2_) were continuously monitored (FUKUDA COLIN Co., Ltd, Tokyo Japan).

The animal was positioned in a stereotaxic holder (Narishige Group, Tokyo Japan), and the headpost was surgically mounted using electrical scalpel (electrosurgical unit, vendor: Semco Co., Ltd Japan, OR Meiji Seika Pharma Co., Ltd, Japan), ceramic screws (Rogue Research, Mtl, CA, Canada) and dental bond (Super-Bond C&B, Sun Medical Co.,Ltd, Japan), which were then covered with acrylic resin (UNIFAST 2, GC Co., Tokyo, Japan). Following the surgery, the animals received antibiotics (Victus, DS Pharma Animal Health Co., Ltd, Japan; 5 mg/ml/kg, i.m.) and analgesic (Metilon, Daiichio-Sankyo Co.,Ltd, Japan, 0.4-0.5 ml) for four days.

### 2.2. Pilot Studies

#### 2.2.1. Structural Image Acquisition

For anatomical reference and headpost surgeries, we acquired structural images using a 24-channel coil designed for anesthetized macaque monkeys (Autio et al., 2020). The macaques were sedated with a combination of atropine (0.2 ml, i.m.), dexmedetomidine (3 µg/kg, i.m.), and ketamine (6 mg/kg, i.m.), after which they were intubated with tracheal tubes. Anesthesia was maintained with intravenous dexmedetomidine (3 µg/kg/hr) and 0.6 % isoflurane administered via a calibrated vaporizer with a mixture of air 0.75 L/min and O_2_ 0.1 L/min. Subjects were ventilated (Cato, Drager, Germany), and catheters were inserted into the caudal artery for blood-gas sampling. Rectal temperature was maintained at approximately 38°C using a blanket.

Structural image acquisition protocols (T1w and T2w) have been previously described (Autio et al., 2020). In brief, T1w images were acquired using a 3D MPRAGE sequence with the following parameters: 0.5 mm isotropic, FOV 128×128×112 mm, matrix 256×256 slices per slab 224, coronal orientation, readout direction of inferior (I) to superior (S), phase oversampling 15%, averages 3, TR 2200 ms, TE 2.2 ms, TI 900 ms, FA 8.3, bandwidth 270 Hz/pixel, GRAPPA 2, turbo factor 224 and pre-scan normalization. T2w images were acquired using a 3D SPACE sequence with the following parameters: 0.5 mm isotropic, FOV 128×128×112mm, matrix 256×256, slice per slab 224, coronal orientation, readout direction I to S, phase oversampling 15%, TR 3200 ms, TE 562 ms, bandwidth 723 Hz/pixel, GRAPPA 2, turbo factor 314, echo train length 1201 ms and pre-scan normalization.

After head-post implementation, all structural image acquisitions were performed using the newly developed 24-channel coil optimized for awake imaging. The imaging parameters remained the same as above, except for the orientation was transversal and phase encoding direction RL.

#### 2.2.2. Quantitative Relaxation Mapping

To estimate ferumoxytol content in the brain vascularity, quantitative transverse relaxation time (T_2_*)-mapping was performed before and after each ferumoxytol injection, as well as at the end of each session to assess potential washout of the ferumoxytol contrast agent. T_2_*-maps were generated using a 3D multi-echo gradient-echo sequence with 1.25 mm isotropic resolution and TEs of 2, 6, 10, 14, 18 and 22 ms. Other imaging parameters were FOV 80×80×80 mm, matrix 64×64×64, phase partial Fourier 7/8, bandwidth 610 Hz/pixel, flip-angle 10°, GRAPPA 2 and TR 50 ms. The scanners image reconstruction and multi-echo fitting algorithm were used to generate quantitative T_2_* (=1/R_2_*) images. In rare cases where the contrast agent injection failed, the experiment was either cancelled (for awake macaques), or an additional dose of ferumoxytol was administered (for anesthetized animals) to achieve target T_2_*-value in gray matter.

To determine the Ernst angle for resting-state fMRI, semiquantitative longitudinal relaxation time (T_1_)-maps were obtained using the MP2RAGE sequence (N=2). The imaging parameters were 0.5 mm isotropic, TR 5000 ms, TE 2.2 ms, TI1 600 ms, TI2 2200 ms, FA1 4°, FA2 4°, bandwidth 270 Hz/pixel, GRAPPA 2 with 24 PE reference lines, turbo factor 168, and pre-scan normalization.

#### 2.2.3. Evaluating Accelerated Imaging

We evaluated the efficiency of multiband slice acceleration factor on fMRI tSNR in anesthetized macaque monkeys. In brief, simultaneous slice excitation with increasing multiband factors reduces the TR, resulting in incomplete T_1_ recovery and a consequent decrease in the optimal (Ernst) flip-angle and tSNR. However, since more data volumes can be acquired within a fixed time window, a more informative measure of data quality is the tSNR scaled by the square root of the number of acquired timepoints (Smith et al., 2013). To account for this, tSNR was evaluated over a matched acquisition time (10 minutes) across various multiband factors (1, 2, 3, 4, 5, 6, 7, and 8), with corresponding minimum excitation and refocusing RF-pulse lengths (maintaining constant spectral width), minimum TRs, blood Ernst angles, and a fixed bandwidth. Each imaging slice was visually inspected for potential multiband leakage artefacts.

#### 2.2.4. Optimizing Ferumoxytol Contrast Agent Dose

After practical in-plane GRAPPA and out-of-plane multiband accelerations were determined, we next set to determine optimal amount of ferumoxytol contrast agent in brain vasculature (see Appendix). For this objective, we used ferumoxytol contrast agent concentration (0, 6, 12, 18, and 24 mg/kg) on resting-state fMRI fluctuations in anesthetized macaques (N=3). Between each injection, quantitative T_2_-maps were acquired.

### 2.3. Resting-state fMRI in Awake Monkeys

For CBVw fMRI, gradient safety limitations (Gradient Safety Watch Dog; Siemens) were disabled to allow stronger slew rates and faster *k*-space coverage, enabling a shorter TE. Additionally, the use of GRAPPA acceleration (factor 2) further contributed to shortening TE (= 15 ms), benefiting from the improved parallel imaging performance enabled by the newly developed coil. The imaging parameters were: 1.25 mm isotropic, FOV 90×90 mm, matrix 72×72, number of slices 48, phase partial Fourier 7/8, bandwidth 1654 Hz/pixel, echo spacing 0.73 ms, TR 755 ms, flip-angle 55°, multiband factor 3, multiband LeakBlock kernel, GRAPPA 2 with a 30 s gradient-echo reference scan to reduce physiological noise in the image reconstruction (Polimeni et al., 2016; Vu et al., 2017), and pre-scan normalization. To determine optimal contrast agent dose, CBVw resting-state fMRI pilot experiments were performed in anesthetized macaque monkeys using varying doses (0, 6, 12, 18 and 24 mg/kg) of ferumoxytol (Feraheme, AMAG Pharmaceuticals Inc, MA, USA).

The B_0_ field-map was estimated using a pair of spin-echo EPI images with opposite phase encoding directions (Andersson et al., 2003) (LR and RL directions, 1.25 mm isotropic, TE 46 ms, TR 4.4 s, prescan normalization). The spin-echo EPI scans were matched with geometry and distortion properties of fMRI acquisitions (FOV 90×90 mm, matrix 72×72, number of slices 48, phase partial Fourier 7/8, bandwidth 1654 Hz/pixel, echo spacing 0.73 ms, GRAPPA 2, fat-suppression, flip-angle 90°, averages 4).

For functional imaging of awake macaques, the subject’s head was securely fixed to the coil using a headpost, and the monkeys were trained to sit in a head-first-sphinx position in an MRI-compatible monkey chair (Fig. 1E, F). The monkeys were gradually accustomed to the auditory noise of the MR pulse sequences (e.g. echo-planar-imaging) and the mock-scanner environment (by MO). Once the training was successfully completed, functional image acquisition was performed using a protocol similar to the CBVw fMRI protocol described above, with the slice orientation set to the coronal direction in scanner coordinates (corresponding to the axial direction in brain coordinates). Each scan session consisted of four single-run scans, each lasting 10 minutes, resulting in a total of 3,116 frames and an overall acquisition time of 40 minutes. Monkeys exhibited no signs of discomfort during the imaging sessions. On the contrary, monkeys had a tendency to become drowsy during imaging sessions. To help maintain arousal levels, subjects were rewarded with apple puree between the 10-min fMRI runs. Additionally, an Inscapes video (www.headspacestudios.org), developed with developmental specialists and widely used in pediatric and adult studies, was displayed via an MRI-compatible monitor (BOLDscreen, Cambridge Research Systems Ltd, Kent, UK) to enhance compliance and reduce head movement and drowsiness (Vanderwal et al., 2015, 2019). The Inscape evokes FC patterns more similar to rest than conventional movies (Vanderwal et al., 2015).

To further monitor alertness, we continuously tracked the animal’s eyes using an MRI-compatible camera. Although our study was not designed to rigorously quantify alertness levels, we estimate that animals appeared drowsy in approximately 5% of scans, in which case we used the scanner’s loudspeaker to alert them. In fewer than 10% of scans, animals had their eyes closed, though this alone does not necessarily indicate drowsiness. No imaging sessions were excluded due to sleep or excessive motion.

Each imaging session was separated by at least by a week to allow washout of ferumoxytol contrast agent. The imaging protocols are publicly available (https://brainminds-beyond.riken.jp/hcp-nhp-protocol).

### 2.4. Data Analysis

#### 2.4.1. Adapting NHP-HCP Data Analysis Pipelines for CBV-Weighted Resting-State fMRI

Data analysis was performed using a non-human primate (NHP) adaptation of the HCP pipelines (NHP-HCP) (Autio et al., 2020; Glasser et al., 2013; Hayashi et al., 2021), which includes both structural and functional preprocessing. The NHP-HCP pipelines utilize FMRB’s Software Library (FSL) version 6.0 (Jenkinson et al., 2012; Smith et al., 2004), Freesurfer version 6 (Fischl, 2012) and Connectome Workbench version 2.0. These pipelines are available at https://github.com/Washington-University/NHPPipelines, and are planned for integration into the HCP pipeline (https://github.com/Washington-University/HCPpipelines). The modifications for CBVw fMRI preprocessing are summarized in Figure 2.

**Figure 2.**
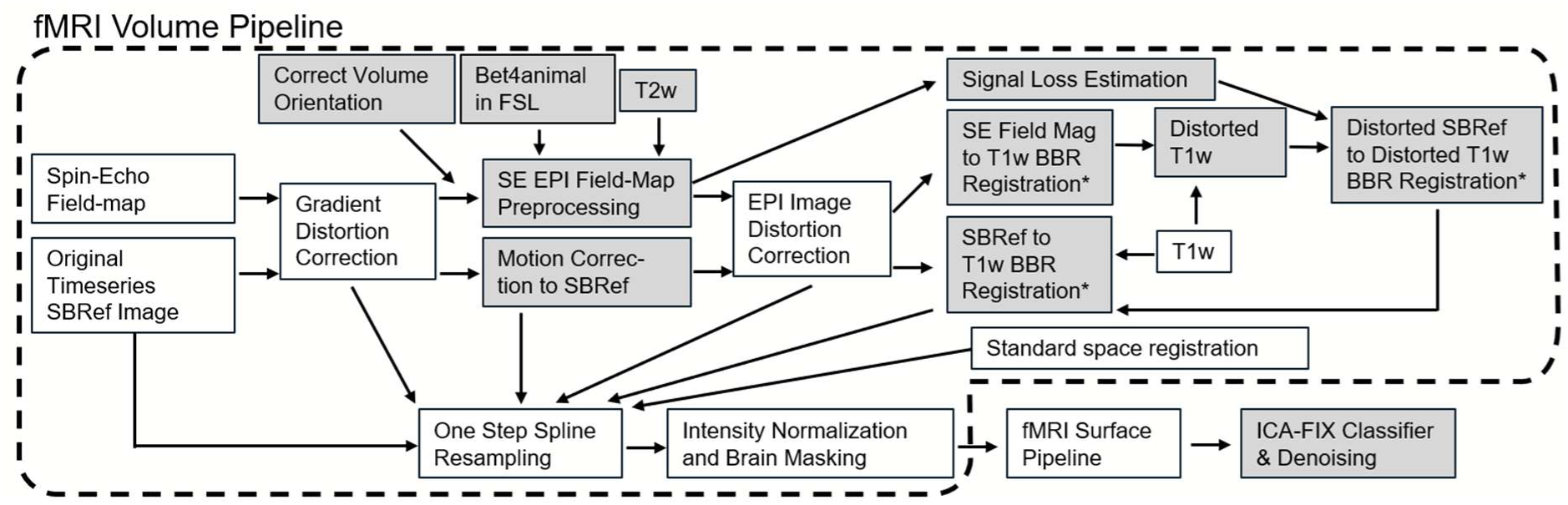
Functional MRI data analysis flow chart. Grey boxes indicate modifications to the previous fMRI pipeline (Glasser et al., 2013; Autio et al., 2020). *BOLD fMRI uses white/grey boundary while CBVw fMRI uses brain outer surface boundary for T1w BBR registration. Abbreviations: BBR: Boundary-based registration; EPI: Echo-planar imaging; ICA: Independent component analysis; LR/RL: Left-Right phase encoding directions; MSM: Multimodal Surface Matching; SBRRef: Single-band Reference scan; and SE: Spin-echo.

#### 2.4.2. Structural Image Pre-Processing

Structural image processing followed our previous protocol and is described in more detail elsewhere (Autio et al., 2020; Glasser et al., 2013). In brief, the NHP-HCP PreFreeSurfer pipeline was used to align structural T1w and T2w images into an anterior-posterior commissural (AC-PC) line using a rigid body transformation, remove non-brain structures, align T2w and T1w images using boundary-based registration (Greve & Fischl, 2009), and correct for signal intensity inhomogeneity. Sphinx-positioned brains, which are often misoriented by the scanner, were corrected using a customized ‘CorrectVolumeOrientation’ script. The removal of non-brain structures and alignment to AC-PC lines were also significantly improved by implementation of ‘bet4animal’ in FSL 6.0.6.6.

FreeSurferNHP pipeline was used to reconstruct the cortical surfaces using FreeSurfer v6.0 (Fischl, 2012). This process included scaling of brain size by a factor of two (using ‘ScaleVolume.sh’), intensity correction, segmentation of the brain into cortex and subcortical structures, reconstruction of the white and pial surfaces and estimation of cortical folding maps and thickness. The intensity correction was performed using FMRIB’s Automated Segmentation Tool (FAST) (Zhang et al., 2001) and whole brain intensity was scaled by a species-specific factor (80). The white matter segmentation was modified by filling a white matter skeleton to accurately estimate white surface around the blade-like thin white matter particularly in the anterior temporal and occipital lobe (Autio et al., 2020). Next, the pial surface was initially estimated by using intensity normalized T1w image and then modified using the T2w image to exclude dura and blood vessels (Autio et al., 2020; Glasser et al., 2013). After this, all the volumes and surfaces were rescaled to the original brain size. The quality of surfaces was visually inspected, and where needed, manual edits were made to the brain, white and ribbon segments (by JA, TI, TH). The surfaces were then re-evaluated by running a pipeline using the edited segments.

The PostFreeSurfer pipeline transformed anatomical volumes and cortical surfaces into the standard macaque space (Hayashi et al., 2021), performing surface registration using folding information via MSMSulc (Robinson et al., 2018). Surface models and data were resampled to a high-resolution 164k mesh (per hemisphere), as well as lower resolution mesh (10k) for processing functional MRI data.

#### 2.4.3. Quantitative Relaxation Mapping

The quantitative T_1_ and T_2_* images were brain masked, aligned into an anterior-posterior commissural (AC-PC) line using a rigid body transformation. Cortical surface mapping (in the 10k CIFTI format) was performed with wb_command –volume-to-surface-mapping using the value of enclosing voxel at the white and gray matter surface. The estimated T_1_ median in gray matter (T_1_=1.37 s) was used to determine Ernst angle for CBVw fMRI (TR 0.75 s yields FA=55°) whereas estimated T_2_* median in grey matter was used to compare optimal TE for CBVw fMRI (TE=ΔT_2_*).

#### 2.4.4. Functional Data Pre-Processing

The CBVw functional image processing was adapted from our previous BOLD NHP-HCP fMRI pipeline (Autio et al., 2020; Glasser et al., 2013), with modifications to address unique pre-processing challenges associated with CBVw imaging. Functional images were motion corrected using MCFLIRT (Jenkinson et al., 2002). Because MCFLIRT has a set of hard-coded assumptions of registration resolution, the brain size was scaled by a factor of two and rescaled to the original before and after MCFLIRT, respectively. Then images were corrected for geometric B_0_-distortions using spin-echo field-maps and TOPUP (Andersson et al., 2003) with adjusted warping resolution and smoothness.

The injection of ferumoxytol contrast agent reduced the signal intensity and contrast between white and gray matter, degrading the alignment of functional images to AC-PC aligned anatomical T1w images. To address this limitation, we implemented a multi-step registration strategy: registration was initialized using the spin-echo field-map magnitude image. This process involved creating a synthetic B_0_-distorted T1w image derived from a B_0_ field-map, which facilitated initial alignment. Subsequent registration was refined using the undistorted spin-echo magnitude image. We applied ‘boundary-based registration’ (BBR) (Greve & Fischl, 2009) to achieve precise alignment. This method used the contrast between the outer brain boundary and cerebrospinal fluid (CSF), using a brain mask in FSL’s BBR and pial surfaces in FreeSurfer’s BBR. The BBR approach effectively weighed the contrast at the brain boundary, resulting in robust and accurate alignment between ferumoxytol fMRI and anatomical images. Initial registration with spin-echo (field-map) magnitude images resulted in low minimum cost function values (0.03 ± 0.01 for FSL’s BBR; 0.42 ± 0.10 for FreeSurfer BBR; n = 108 sessions). Single-band gradient-echo registration also yielded low values (0.11 ± 0.02, max = 0.16 for FSL’s BBR; 0.08 ± 0.04, max = 0.25 for FreeSurfer BBR). BBR was performed using T1w contrast (slope=0.5 in FSL’s BBR). For FSL’s BBR, a brain mask volume was created from pial surfaces using the FreeSurferNHP pipeline. FreeSurfer BBR employed a white matter projection absolute distance of 0.7 mm to account for the thin, plate-like white matter in the macaque brain. Finally, functional data were normalized to the grand 4D mean and masked to include gray matter structures.

The cerebral cortical gray matter voxels were mapped to the surface with the partial-volume weighted ribbon-constrained volume to surface mapping algorithm and voxels having large deviations (greater than 0.5 s.d. above mean) from the local (in a 5 mm sigma Gaussian) neighborhood voxels’ coefficient of variation were excluded (Glasser et al., 2013). Data was minimally smoothed at 1.25 mm full-width half-maximum (FWHM) using geodesic Gaussian surface smoothing algorithm with vertex area correction and resampled according to the folding-based registration (MSMSulc) to a standard mesh in which the vertex numbers correspond to neuroanatomically matched locations across subjects. The subcortical gray matter voxels were processed in the volume using 1.25 mm FWHM subcortical parcel-constrained smoothing and resampling. Altogether, these processes transformed the functional data into a standard set of greyordinates (≈10,000 [10k] vertices per hemisphere and ≈8,000 [8k] subcortical voxels) in the Connectivity Informatics Technology Initiative (CIFTI) format (Autio et al., 2020; Glasser et al., 2013).

#### 2.4.5. Group-Level Functional Connectivity

Structured temporal noise, which arises from imaging artifacts such as motion and physiological fluctuations, was identified and removed using a macaque-adapted version of FMRIB’s spatial ICA-based X-noisefier (FIX) (“ICA + FIX”) (Glasser et al., 2018; Griffanti et al., 2014, 2017; Salimi-Khorshidi et al., 2014). To further distinguish between meaningful neural signal components from unstructured noise, we applied principal component analysis (PCA). This allowed us to separate data into two subspaces: (1) a structured subspace, which contains signal that exhibit meaningful spatial and temporal patterns, and (2) an unstructured subspace, which consists of signal components that follow a random statistical distribution (modelled as a Wishart null distribution) and are therefore considered noise (Glasser et al., 2016). Unstructured (Gaussian) noise primarily originates from the MRI system itself, including the RF-transmit coil, the same (head), the gradient coils, the RF-receive coil, and pre-amplifier. In contrast, structured (non-Gaussian) noise results from imaging artifacts such as motion and physiological fluctuations, which exhibit characteristic spatial and temporal patterns. The ICA-FIX cleaned structured subspace was further analyzed at the group-level functional connectivity by correlation of each vertex’s eigenvalues with each other and computing a dense Pearson’s correlation matrix.

Structured temporal noise arising from imaging artifacts (e.g. motion and physiological noise) were manually classified and removed using a macaque-adapted version of FMRIB’s ICA based X-noisefier (FIX) (“sICA + FIX”) (Glasser et al., 2018; Griffanti et al., 2014, 2017; Salimi-Khorshidi et al., 2014). Principal component analysis (PCA) was applied to segregate data into structured and unstructured sub-spaces (the latter following Wishart null distribution) (Glasser et al., 2016), and the structured subspace was decomposed into statistically independent components using spatial ICA. The resulting components were manually classified as “signal” or “noise”, based on their spatial distribution and temporal properties (Griffanti et al., 2014).

The CBVw FIX classifier was then trained on this manual classification (N=3; total trials=27; total data points=84,132) and its performance was characterized by true positive rate and true negative rate. Classifier performance was evaluated using leave-one-out accuracy testing across a range of thresholds. The training file will be included in the coming FSL release.

Information on the different categories of CBVw fMRI fluctuations was evaluated using the HCP’s RestingStateStats adapted for macaque monkeys (Autio et al., 2020; Marcus et al., 2013). In brief, RestingStateStats script decomposes total CIFTI fMRI timeseries variance (prior to any preprocessing) into six categories: low frequency noise (by high-pass filter), motion, artifacts and nuisance signals (by FIX classification), unstructured noise (by PCA, see above), neural CBVw fluctuations (by FIX classification), and FIX-cleaned mean greyordinate timeseries (MGT). The fractional contribution of each category was calculated by dividing by the total fMRI variance in the grayordinate CIFTI format, which contains cortical gray matter in surface GIFTI format and subcortical structures in volume NIFTI format. The CNR was determined as the ratio of neural variance to unstructured noise variance.

To investigate within- and between-subjects FC reproducibility, the cleaned fMRI timeseries from each imaging session were parcellated using the M132 atlas (Markov et al., 2014). A Pearson’s correlation matrix was calculated to represent FC for each session. The lower triangular portion of each session’s matrix was compared across sessions using Spearman’s rank correlation coefficient (Autio et al., 2021).

Subject identification accuracy was quantified by identifying the session pair with the highest Spearman’s rank coefficient. A score of 1 was assigned if the highest correlation corresponded to the same subject, and a score of 0 was assigned otherwise. Scores were averaged across all sessions to compute overall identification accuracy. Additional subject identification analysis was performed using noise-dominant timeseries, in which neural components were regressed out and the noise components were kept in the timeseries.

To explore the functional organization of the macaque brain using group-level FC analyses, we first normalized each session’s dense timeseries for variance to account for potential differences in ferumoxytol concentrations across sessions and subjects. We then re-constructed the group data using a Wishart null distribution threshold to remove redundant unstructured noise in the eigenvalue subspace (Glasser et al., 2016). The reconstructed data was subsequently used to compute a Pearson’s correlation matrix and examine FC across the macaque gray matter.

## 3. Results

### 3.1. Coil Performance

Table 1 provides an overview of the quality assurance measures conducted on the newly developed 24-channel phased-array coil for awake macaque imaging. Our assessment revealed low coupling between adjacent coil elements and low noise correlation.In-plane acceleration using GRAPPA R=2 resulted in an inverse g-factor marginally above unity, followed by slight reduction in R=3, and a notable drop at R=4 reflecting the upper limit of the coil’s parallel imaging capability (Fig. 3C; Supp. Fig. 2).

**Table 1.** Quality assurance of awake (top row) and anesthetized (bottom row) macaque 24-channel phased-array receive coils. Q-ratio is the unloaded to loaded ratio of the individual coil elements. Decoupling between adjacent elements indicates low mutual inductance between the elements. This result is supported by low average noise correlation between the coil elements (mean ± std). Inverse geometry (1/g) factor above unity indicates a small noise cancellation during parallel image reconstruction using a reduction factor R=2. Relatively homogeneous flip-angle distribution over cortical surface indicates that the coil did not interfere with B_1_-transmission.

| Coil | Q-ratio | Decoupling (dB) | Noise corr. | 1/g | $B_1+$ | Reference |
| --- | --- | --- | --- | --- | --- | --- |
| Awake | 215/75<br>=2.9 | -20 | 0.08 ±<br>0.05 | 1.14 ±<br>0.11 | 89.0° ±<br>4.5 | |
| Anesthetized | 215/75<br>=2.9 | -20 | 0.08 ±<br>0.04 | 1.03 ±<br>0.07 | 86.6° ±<br>2.3 | (Autio et al., 2020) |

**Figure 3.**
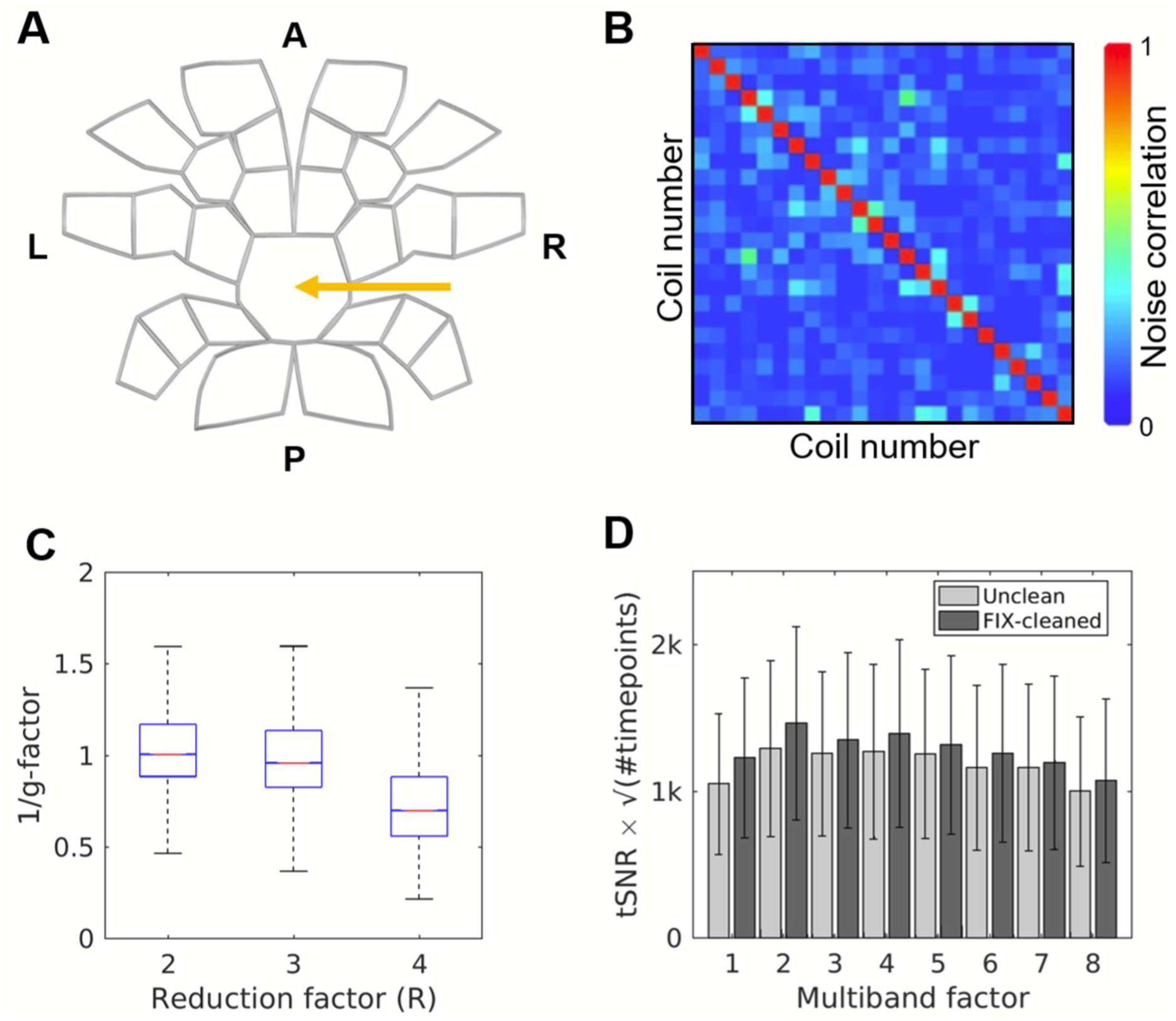
Assessment of the 24-channel coil’s parallel imaging performance. **(A)** Flattened map of the coil element arrangement. The orange arrow indicates the opening in the phased-array designed to accommodate the rigid head attachment. **(B)** Noise correlation matrix. **(C)** Inverse geometry (1/g)-factor maps obtained using gradient-echo imaging with generalized autocalibrating partially parallel acquisitions (GRAPPA) at different reduction factors. **(D)** Multiband performance evaluation assessed using the temporal signal-to-noise ratio (tSNR) scaled by square root of the number of timepoints, across different multiband factors.

We also evaluated the coil’s performance for multiband acceleration and observed an increase in tSNR (scaled by the square root of timepoints) at moderate acceleration factors (2-4) followed by a gradual decline (Fig. 3D), along with notable cross-slice artefacts (data not shown). These quality assurance measures closely resembled those obtained with our previous 24-channel phased-array coil tailored for imaging anesthetized monkeys (Autio et al., 2020), with the only notable difference being a slightly elevated variation in the B_1_+ transmission field attributable to the coil’s positioning 10 cm above the B_0_ isocenter in the awake configuration. Collectively, these quality assurance results indicate robust parallel imaging capabilities, which are essential for accelerated CBVw fMRI in awake macaque monkeys.

Representative structural brain images acquired with parallel imaging acceleration are shown in Figure 4. The image quality is comparable to that obtained with our previous coil for anesthetized animals, (Fig. 4A, B), although the newly developed coil exhibits slightly lower SNR. Some signal dropout is inevitable due to the phased-array opening designed to accommodate head-post fixation. Signal loss is observed near the top of the brain, where the B_0_ receive field is attenuated relative to our previous coil (Fig. 4C, D). Despite the B_0_ biasfield variations of approximately 50% throughout the cortex, SNR remains sufficient to enable robust image segmentation and accurate delineation of white matter and pial surfaces (Fig. 4A, red and yellow lines).

**Figure 4.**
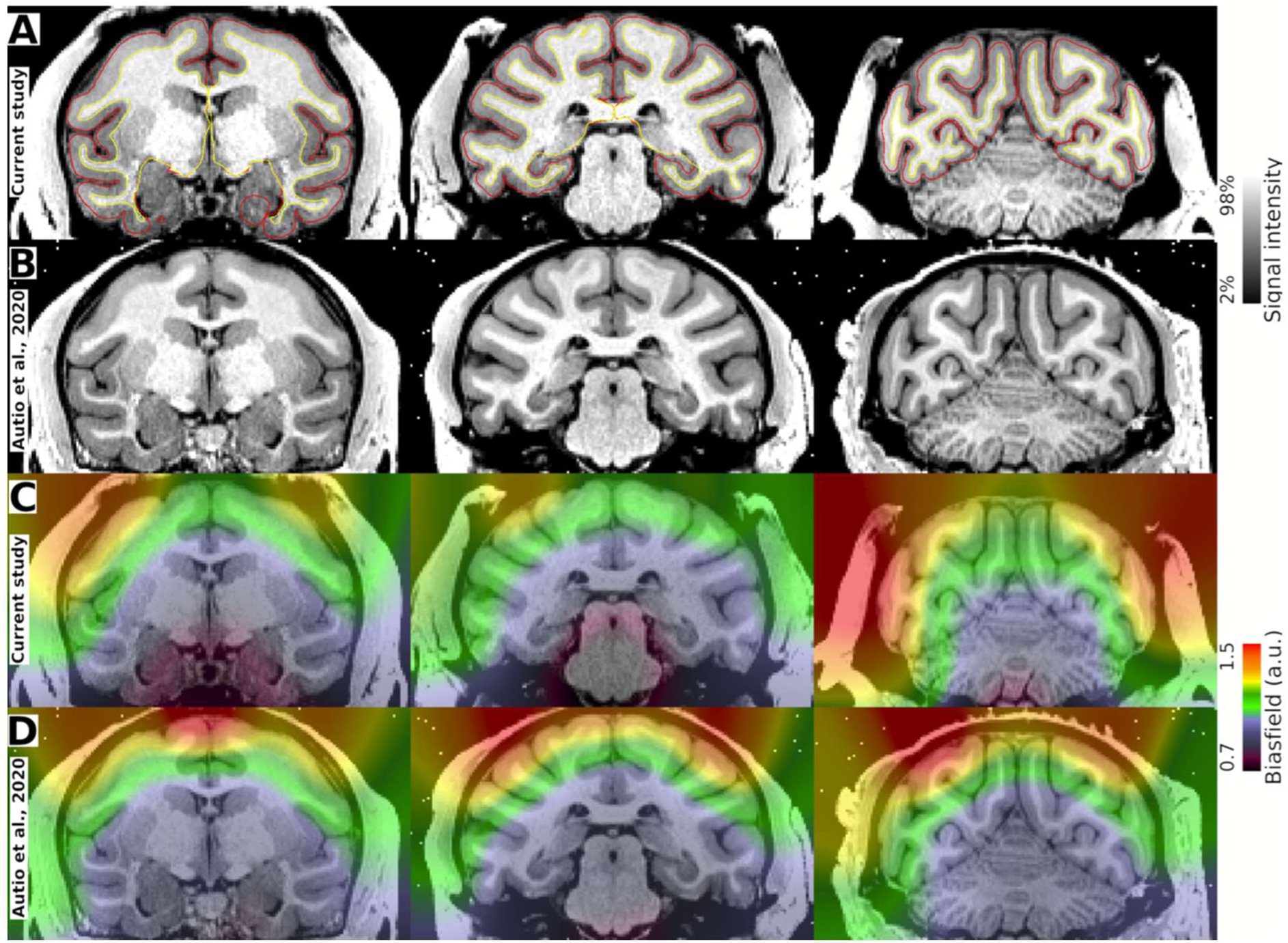
Example of structural image quality assessment. Structural T1w images acquired using a phased-array coil designed for **(A)** awake and **(B)** anesthetized monkeys (Autio et al., 2020). The image resolution is 0.5 mm isotropic, GRAPPA 2, and acquisition time 17 min. The red and yellow lines indicate pial and white matter surfaces, respectively. **(C, D)** B_0_ biasfield estimated from T1w and T2w images.

### 3.2. Contrast Agent Dose Optimization

Pilot experiments were conducted to evaluate the impact of ferumoxytol contrast agent concentration (0, 6, 12, 18, and 24 mg/kg) on resting-state fMRI fluctuations in anesthetized macaques (N=3). The objective was to quantify the relative variance of unstructured noise, low-frequency noise filtered noise, structured noise, neural signal fluctuations, and FIX-denoised mean global timeseries (MGT) (Autio et al., 2020; Marcus et al., 2013). The relative neural signal variance exhibited a dose-dependent response to ferumoxytol contrast agent, peaking at concentrations between 12 and 18 mg/kg, and declining at 24 mg/kg (Fig. 5A, B). These results align with classical single-tissue-compartment CNR optimization principles, where the TE (15 ms) closely matches the average T_2_* of gray matter at approximately 16 ms and 14 ms for doses of 12 mg/kg and 18 mg/kg, respectively (see Appendices, Data Acquisition Strategy) (Boxerman et al., 1995; Mandeville et al., 2004). Considering the marginal difference in CNR between doses the 12 and 18 mg/kg doses and the sufficient tSNR across the gray matter (Fig. 5C), the 12 mg/kg dose was chosen for subsequent awake CBV fMRI experiments.

**Figure 5.**
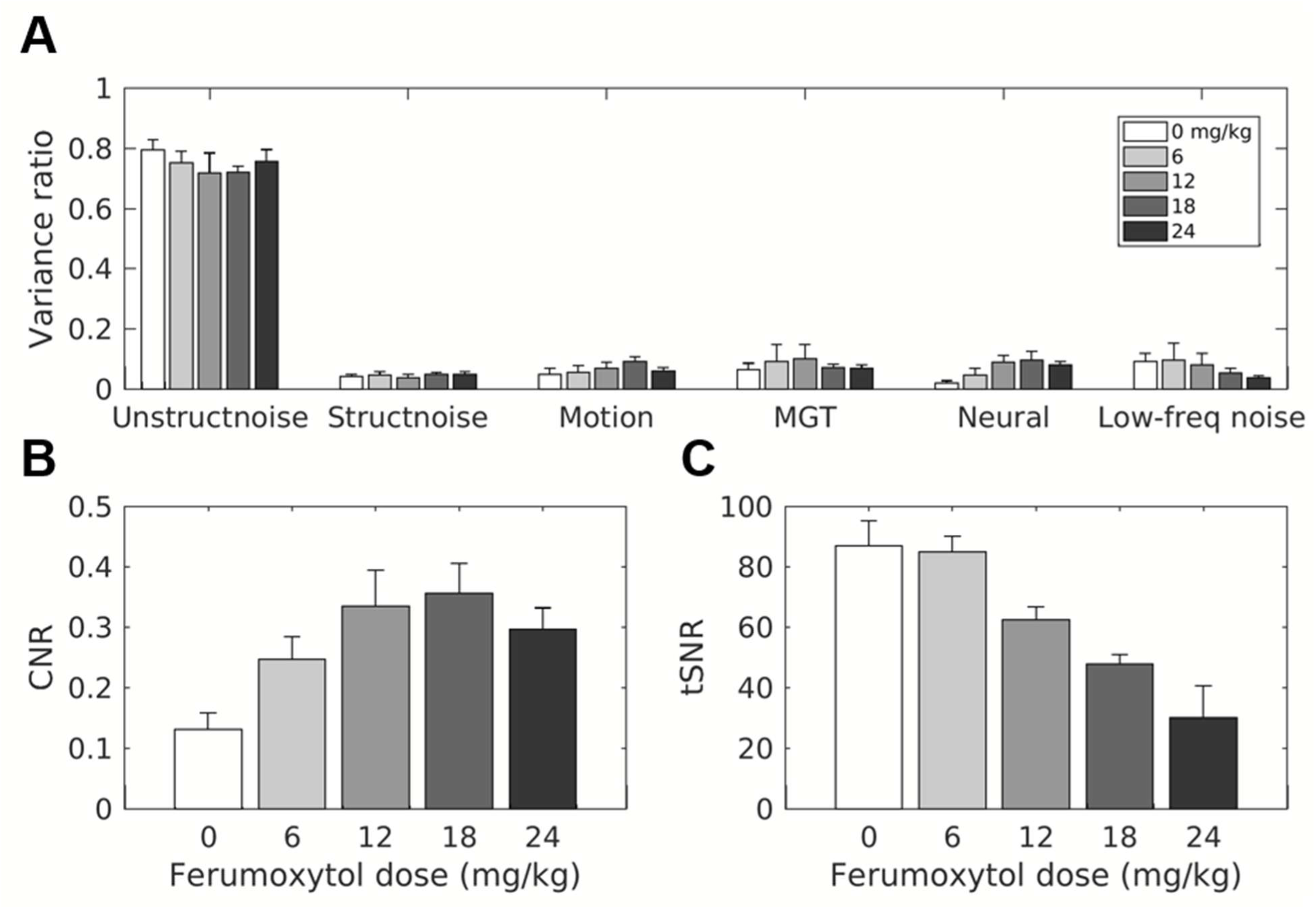
Determining optimal ferumoxytol contrast agent dose for accelerated cerebral blood volume weighted (CBVw) resting-state fMRI. **(A)** Variance categories, **(B)** contrast-to-noise ratio (CNR), and **(C)** temporal signal-to-noise ratio (tSNR) at varying ferumoxytol doses. Data represent mean, with error-bars indicating the standard deviation across image runs and subjects (n = 2 × 3 = 6). Data was analyzed using the Human Connectome Project’s RestingStateStats (Autio et al., 2020; Marcus et al., 2013). Abbreviations: Low-freq noise, low-frequency noise (by high-pass filter); StructNoise, structured noise (by FIX classification); Signal (by FIX classification); UnStruct Noise, unstructured noise (by PCA), MGT FIX-cleaned mean greyordinate timeseries.

It is also noteworthy that tSNR, often used as a quality index in fMRI (with high tSNR indicating good data and low tSNR poor data), gradually decreases as ferumoxytol contrast agent concentration increases (Fig. 5C). This occurs because ferumoxytol injection reduces overall SNR white amplifying neural signal fluctuations (Fig. 5A, B), both leading to a weak tSNR, which is defined as tSNR = signal fluctuation / mean MR signal. For this reason, we consciously refrained from relying on tSNR maps, and instead focused on quantifying neural signal content, as identified using spatial ICA.

### 3.3. ICA-FIX Cleanup and Reproducibility of Functional Connectivity

To evaluate the fMRI data quality, we categorized fMRI timeseries timeseries (totaling ≈1,000 min and ≈84,132 volumes) into unstructured noise (via PCA), structured noise (via spatial ICA) motion, mean grayordinate timeseries, neural signal (via ICA) and low-frequency noise using the FIX (Griffanti et al., 2014, 2017; Salimi-Khorshidi et al., 2014) using a modified HCP resting-state analysis pipeline for macaque monkeys (Autio et al., 2020; Marcus et al., 2013). The average spatial distribution of each category is shown in Figure 6A.

**Figure 6.**
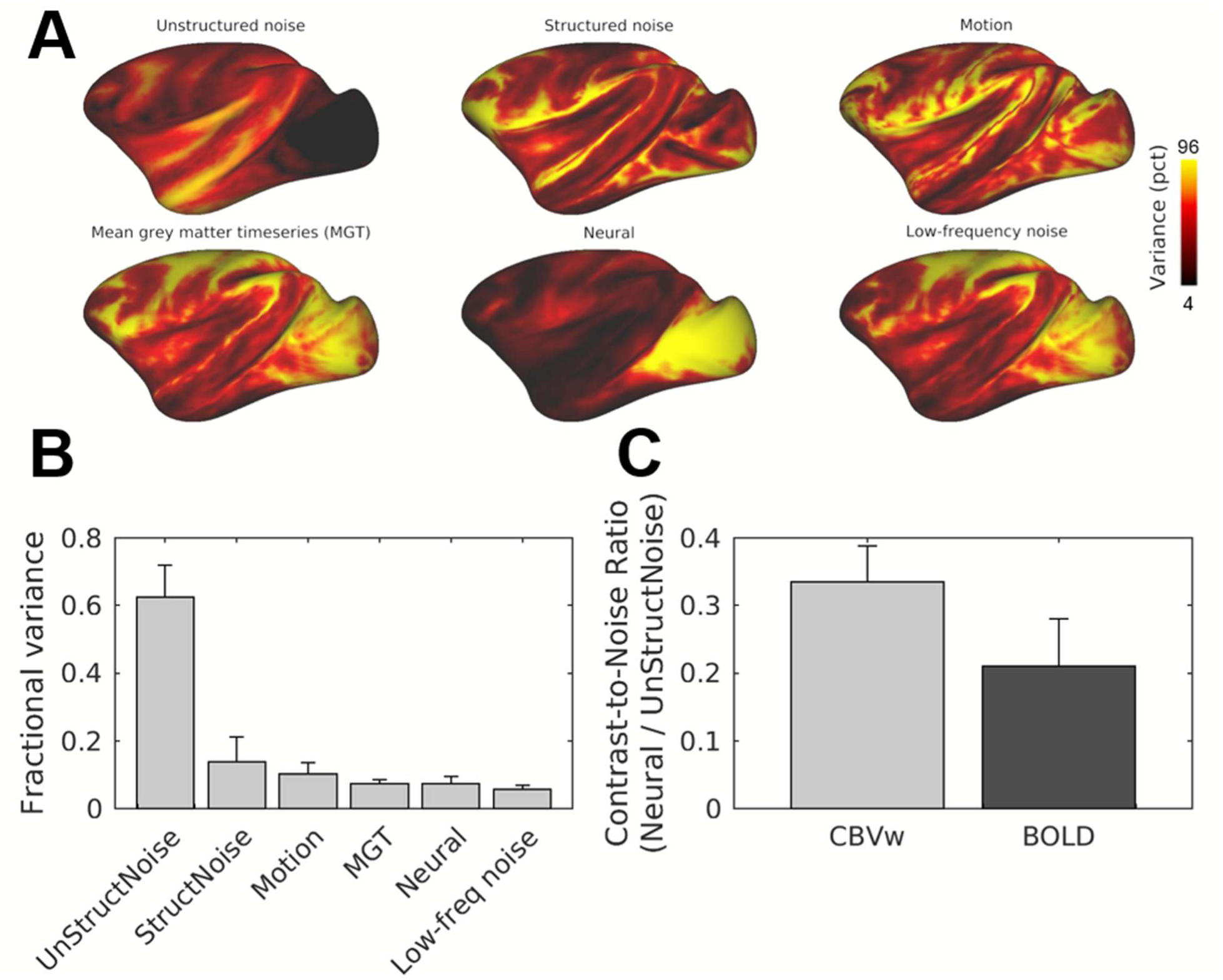
Classification of Noise and Neural Variance in Resting-State fMRI Timeseries. **(A)** Averaged resting-state categories displayed on a cortical flat-map (N=3, total n=27 sessions). **(B)** Summary of average fractional variances across the categorized signals. **(C)** Comparison of contrast-to-noise ratio (CNR) between cerebral awake blood volume -weighted (CBVw) and anesthetized blood oxygen level dependent (BOLD) resting-state fMRI, calculated as the ratio of neural signal variance to unstructured noise variance ratio. Fractional variances for each category were estimated using the HCP RestingStateStats pipeline (Autio et al., 2020; Marcus et al., 2013). Abbreviations: UnStructNoise: UnStructured noise; StructNoise: Structured noise; MGT: Mean gray matter timeseries.

On average, unstructured noise accounted for the largest portion of fMRI signal variance (60%) (Fig. 6B). Unstructured noise was higher in brain regions more distant to coil elements, as expected. In contrast, neural signals were strongest in the visual cortex, likely due to its proximity to coil elements as well as its high baseline vascular volume and neuron density (Autio et al., 2025; Collins et al., 2010). Structured noise, motion and low-frequency noise in total contributed to a 4-fold larger amount of variance than neural signals (30% vs 7%). After removing these artifacts, the resulting CNR, defined as the ratio between neural signal and unstructured noise variances, was significantly larger than in anesthetized BOLD fMRI (p < 10^-7^, Fig. 6C) (Autio et al., 2020).

An exemplar FC map seeded from bottom of the anterior sulcus (Fig. 7A) shows pronounced differences before and after ICA-FIX cleanup (Fig. 7C, D). Correlating all seed-based FCs before and after FIX cleanup reveals that FC is more profoundly improved by FIX in the anterior regions of the brain, with minimal impact in the posterior visual system (Fig. 7B). This anterior-posterior gradient can be partially attributed to the head-post fixation at the posterior part of the skull (Fig. 1), which likely results in the anterior regions of the brain being more susceptible to subtle motion artifacts compared to the posterior regions (Fig. 6A; see motion panel).

**Figure 7.**
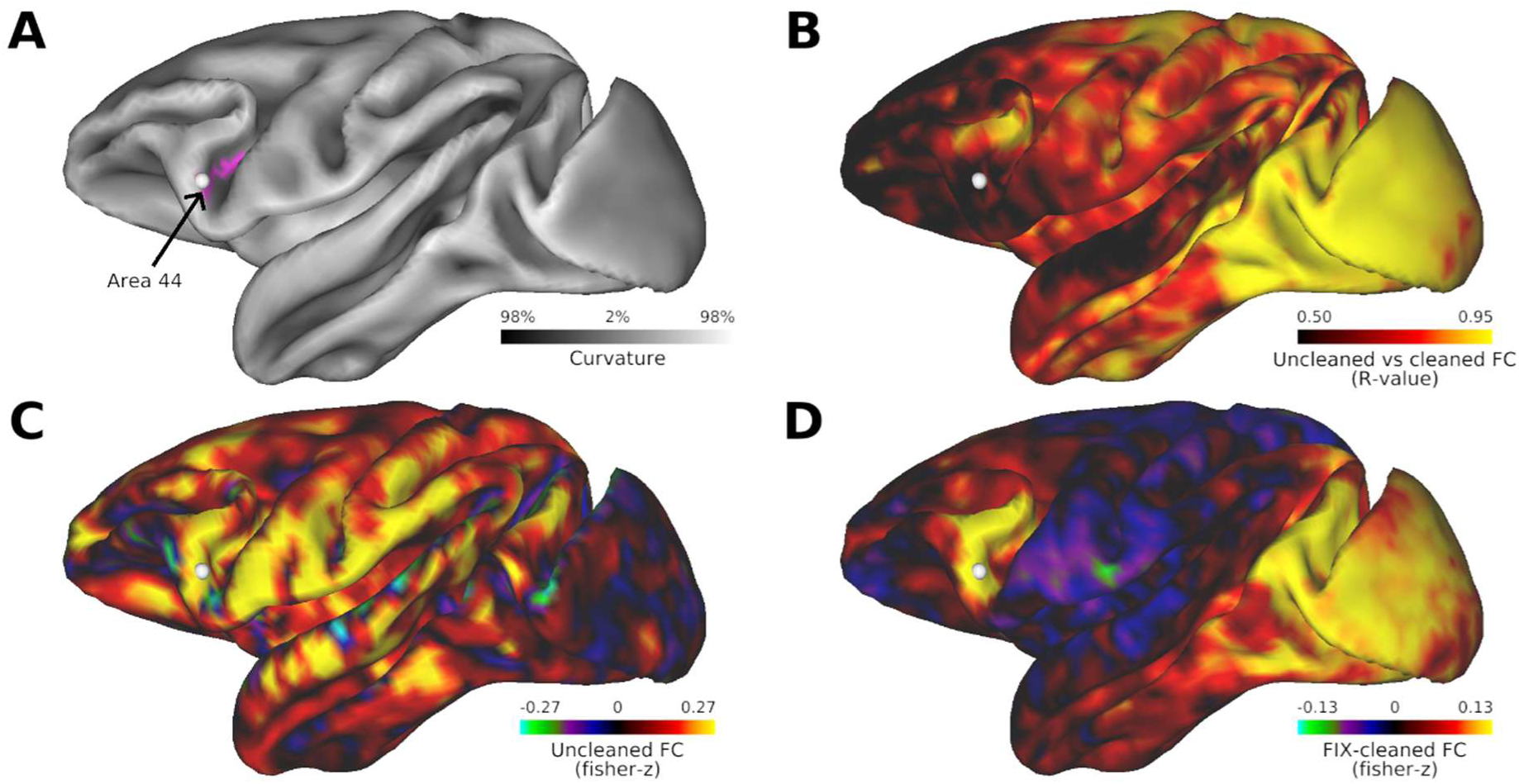
Accelerated Ferumoxytol-weighted fMRI in Combination ICA-FIX Artifact Cleanup Improves Functional Connectivity Estimation in Awake Macaque Cerebral Cortex. **(A)** Seed location at area 44 (white circle) as indicated by M132-atlas (magenta). **(B)** Spatial correlation patterns between FIX-uncleaned and FIX-cleaned functional connectivity (FC) (N=1; imaging sessions=9). Note that the correlation between FIX-uncleaned and cleaned FC varied between 0.53 - 0.95 (5th and 95th percentile), highlighting the importance of data-driven ICA de-noising in ferumoxytol-weighted fMRI. Exemplar seed FC from area 45 **(C)** prior to and **(D)** after FIX-cleanup.

Manually classified sICA components (on average 87 ± 53 noise and 18 ± 7 signal components per fMRI session) were used to train an automatic classifier (python FIX; pyFIX) for denoising ferumoxytol-weighted fMRI data. The classifier demonstrated excellent performance (Table 2), with leave-one-out testing showing a perfect true positive rate of 100% ± 0%. True negative rate ranged from 85% to 100%, depending on the threshold level. Consequently, a threshold of 40 was selected for automatic classification, resulting in a mean true positive rate and true positive rate of 100%.

**Table 2.** FIX classification accuracy test by leave-one-out (LOO) in 27 imaging sessions of awake macaque data. Abbreviations: TPR: true positive rate of signal components; TNR: true negative rate of true artifact components.

| FIX threshold | 1 | 2 | 5 | 10 | 20 | 30 | 40 | 50 |
| --- | --- | --- | --- | --- | --- | --- | --- | --- |
| TPR (mean) | 1.00 | 1.00 | 1.00 | 1.00 | 1.00 | 1.00 | 1.00 | 1.00 |
| TNR (mean) | 0.86 | 0.90 | 0.95 | 0.97 | 0.98 | 0.99 | 1.00 | 1.00 |
| TPR (median) | 1.00 | 1.00 | 1.00 | 1.00 | 1.00 | 1.00 | 1.00 | 1.00 |
| TNR (median) | 0.85 | 0.92 | 0.97 | 0.98 | 0.99 | 1.00 | 1.00 | 1.00 |

Next, we investigated the dependence of FC reproducibility on scan duration using M132 atlas-parcellated correlation matrices (Birn et al., 2013; Laumann et al., 2015). We analyzed data from three awake macaques, each with nine imaging sessions (totaling ≈1,000 min and ≈84,132 volumes) and from five anesthetized each with two imaging sessions (totaling 1,020 min and 81,920 volumes). We generated parcel timeseries and performed within-scan reproducibility analysis by systematically varying the number of fMRI volumes (Supp. Fig. 3A). Our analysis revealed a reproducibility of ρ=0.94 ± 0.03 (Spearman rank ± standard deviation) in awake macaques and ρ=0.94 ± 0.04 in anesthetized macaques (Supp. Fig. 3B). Notably lower reproducibility was observed in subcortical structures in anesthetized macaques (data not shown), suggesting an impact of anesthesia on the consistency of connectivity patterns, which could be further explored to understand differences in brain dynamics between these states.

The within-subject reproducibility of the M132 atlas-parcellated FC matrix was excellent ρ=0.84 ± 0.08 (Fig. 8A, B). The between-subject FC matrix correlation was moderate 0.66 ± 0.08, and comparable to those reported in humans (Koike et al., 2021). The lower between-subject correlation indicates that each subject’s FC matrix is distinct. Notably, each individual could be accurately distinguished across all imaging sessions using any random pair of scans, achieving 94% subject identification accuracy. The scans that were not correctly identified had large, persistent motion artifacts (data not shown). This furthers the idea that FC serves as a reliable neural fingerprint to identify individual humans (Finn et al., 2015) also in macaques based on their distinctive neural activity patterns.

**Figure 8.**
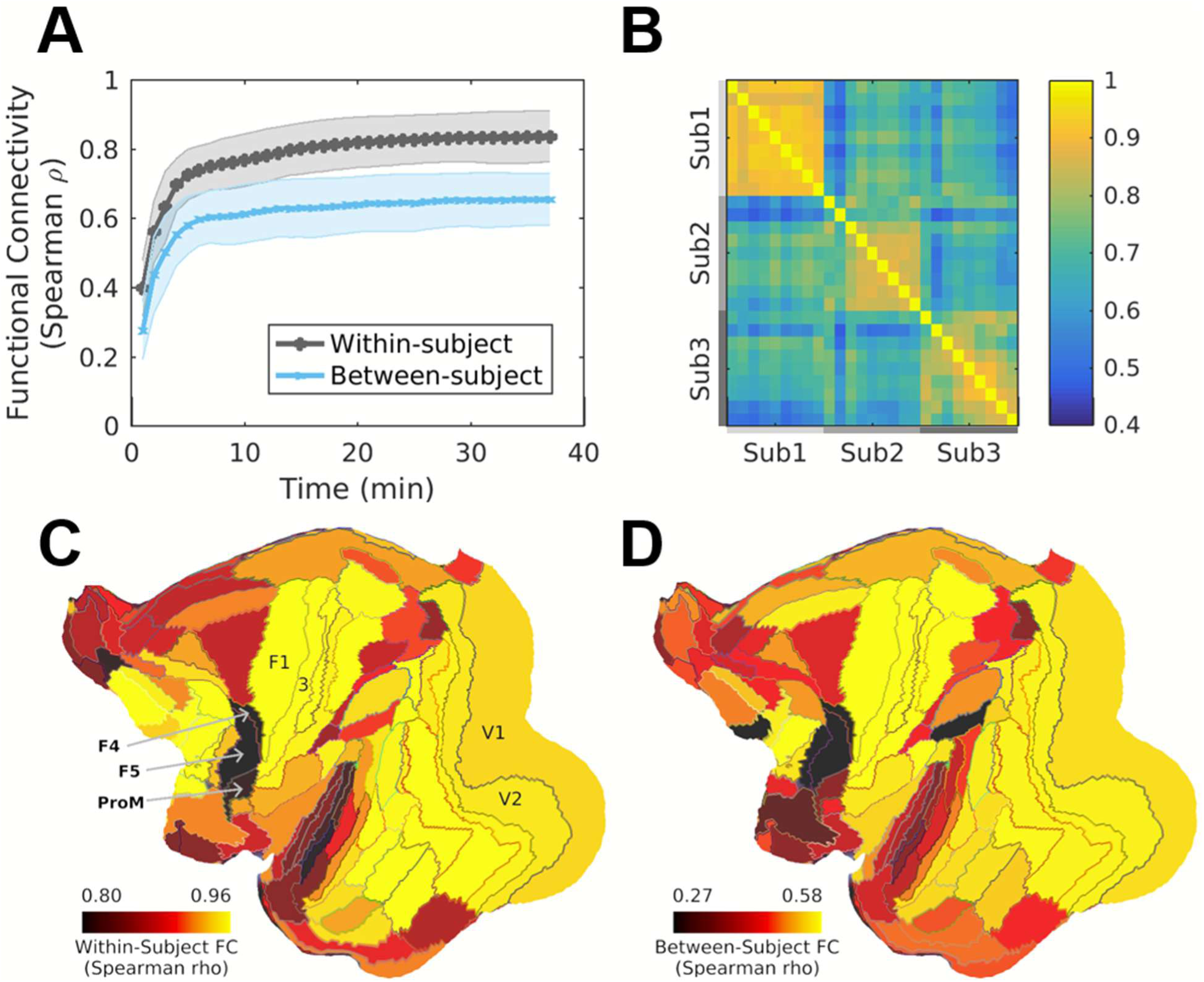
Within- and Between-Subject Reproducibility of Resting-State Functional Connectivity. **(A)** Within-subject and between-subject functional connectivity (FC), parcellated using the M132 atlas, plotted as a function of scan duration. The continuous line represents the mean Spearman’s rank correlation, while the shaded error-bars represent the standard deviation. **(B)** Reproducibility of within-subject and between-subject FC. **(C)** M132-parcellated FC matrices from three awake macaques across nine imaging sessions each. **(D)** Parcel-wise FC reproducibility within-subjects (left) and between-subjects (right) displayed on a cortical flat-map. A total of 182 parcels were analyzed, yielding 16,380 parcellated FC values across both hemispheres of the cerebral cortex.

To further investigate individual differences, we examined noise patterns (e.g. noise classified ICAs) that might also be subject-specific due to variations in large vessel locations, brain shape, and idiosyncratic movements. The within-subject reproducibility of the M132 atlas-parcellated noise matrix was good (ρ = 0.79 ± 0.07), while the between-subject correlation was moderate (ρ = 0.56 ± 0.07) (Supp. Fig. 4), indicating that each subject’s noise correlation matrix is distinct. These noise patterns alone enabled high identification accuracy across subjects, reaching 98%. This demonstrates that noise-dominant correlation patterns can also serve as reliable identifiers of individual macaques.

### 3.4. Group-level Functional Connectivity

Having established excellent data reproducibility at the individual subject level, we next aimed to explore functional organization of the macaque brain using group-level connectivity analyses. For this objective, we re-constructed the group data using a Wishart null distribution threshold for removing redundant unstructured noise (Glasser et al., 2016). An example of seed-based FC from the upper-limb motor cortex (M1; Fig. 9A) is shown in Figure 9B.

**Figure 9.**
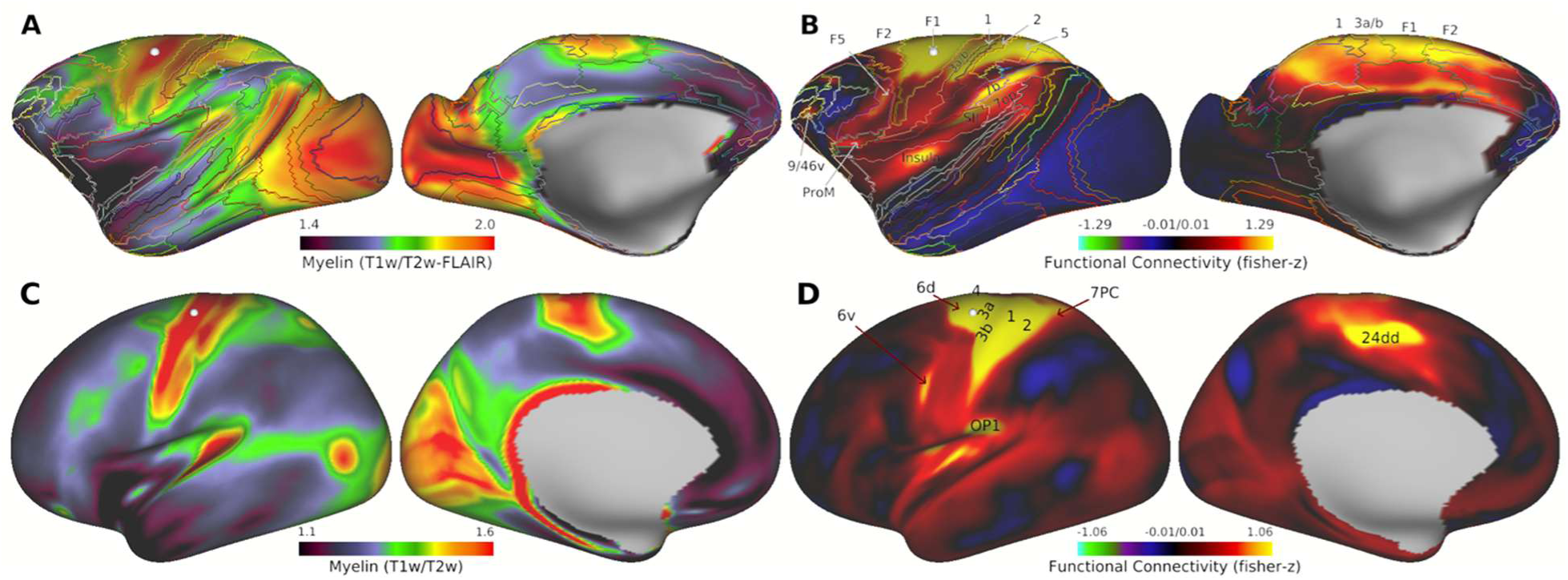
Comparison of Group-Level Resting-State Functional Connectivity in Awake Macaque Monkeys and Humans. **(A)** Macaque myelin map (Autio, et al., 2024). The lines indicate M132-atlas area boundaries (Markov et al., 2014). **(B)** Resting-state functional connectivity (FC; N=3; total n=27) seeded from the primary motor cortex region of upper-limb presentation. **(C, D)** Human myelin map and temporal-ICA cleaned FC (N=1200). Notably, the upper-limb presentation appears to cover a larger fraction of the cerebral cortex in macaque than humans. Abbreviations: M1: primary motor cortex; OP1: Opercular area 1; SII: Secondary somatosensory area; 1: area 1; 2: area 2; 3a: Primary sensory area 3a; 3b: Primary sensory area 3b; 5: area 5; 6a: area 6 anterior; 6d: area 6 dorsal; 6dr: area 6 dorsorostral; 6v: area 6 ventral; 7a: area 7a; 7b: area 7b; 7op: area 7 opercular cortex; 7PC: area 7 of parietal cortex; 9/46d: area 9/46 dorsal; 9/46v: area 9/46v ventral; 24dd: area 24d dorsal.

Seeding from a similar location in human HCP data reveals a strikingly similar spatial pattern when compared to macaques (Fig. 9C, D). The upper limb network in macaques appears to occupy a substantially larger proportion of the cerebral cortex compared to humans. Likewise, seeding from the lateral geniculate nucleus reveals comparable subcortico-cortical FC patterns, though their relative extent varies with respect to the cerebral cortex (Fig. 10A-F). Together, these results demonstrate the effectiveness of the newly developed 24-channel RF receive coil, ferumoxytol-weighted fMRI, and modified NHP-HCP pipelines in mapping the functional organization of the macaque grey matter.

**Figure 10.**
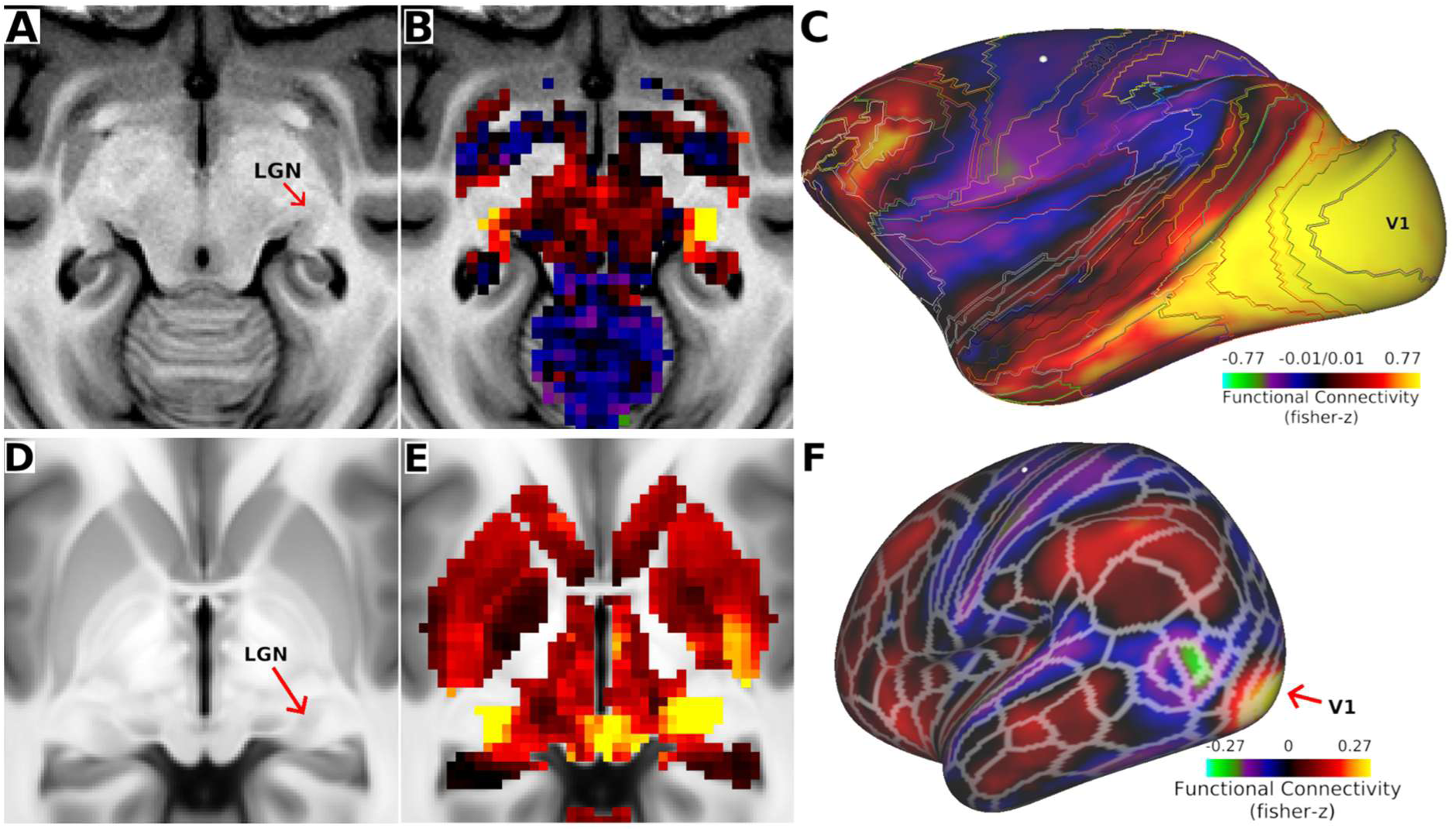
Exemplar Subcortico-Cortical Resting-State Functional Connectivity in Awake Macaque Monkeys and Humans. **(A)** Seed location in the lateral geniculate nucleus (LGN), **(B)** subcortical and **(C)** cortical functional connectivity patterns. **(D, E, F)** Human temporal-ICA cleaned FC (N=1200). Cerebral cortex is rotated to display V1. Abbreviations: V1: Primary visual area.

## 4. Discussion

Here, we have presented a neuroimaging approach for awake macaque monkeys using a combination of a custom-made headpost compatible 24-channel receive RF coil, three-by-two accelerated (out-of-plane MB and in-plane GRAPPA) ferumoxytol-weighted fMRI and modified HCP-NHP preprocessing and analysis pipelines. Importantly, this approach facilitates robust and reproducible functional investigations, effectively addressing the reproducibility challenges associated with NHP neuroimaging (Autio et al., 2021). The resources developed, including the 24-channel macaque coil (via Rogue Research, Montreal, Canada https://www.rogue-research.com/; produced by Takashima Seisakusho Co. Ltd. Tokyo, Japan) along with the data acquisition protocols (https://brainminds-beyond.riken.jp.hcp-nhp-protocol) and NHP-HCP CBVw fMRI analysis pipelines (https://github.com/Washington-University/NHPPipelines), are publicly available. Raw and minimally preprocessed awake ferumoxytol-weighted resting-state fMRI data will also be shared in an upcoming publication, as a part of the NHP-HCP neuroimaging and neuroanatomy project (Hayashi et al., 2021), enabling the broader scientific community can leverage these developments for further research.

The newly developed coil provides good quality structural images important for accurate segmentation and robust cortical surface generation (Fig. 4A), a fundamental requirement for surface-based fMRI analysis. Surface mapping is increasingly important for integrating the growing availability of histological, receptor, genetic, and transcriptomic datasets (BICAN refs; Chen et al., 2023; Froudist-Walsh et al., 2023; Hayashi et al., 2021). However, if high-resolution structural images are required, such as those aiming for cortical layer-specific analysis, we recommend using the 24-channel coil designed for anesthetized macaque monkeys (Autio et al., 2020; 2024), as maintaining awake animals in a prone position under anesthesia can be challenging. Although the benefits of contrast agents have been well-documented in task-based fMRI studies (Leite et al., 2002; Mandeville et al., 2004; Vanduffel et al., 2001; Zhao et al., 2006), the majority of macaque resting-state fMRI research at 3T continues to rely on BOLD contrast (Autio et al., 2021; Milham et al., 2018). However, the relative neural BOLD signal variance in anesthetized macaque monkeys at 3T is modest (≈0.1 - 3%) (Autio et al., 2021; Milham et al., 2018), even when using multiband accelerated imaging sequences (≈4%) (Autio et al., 2020). Ferumoxytol contrast agents provide means to amplify the resting-state fMRI signal fluctuations, increasing the relative signal variance to approximately 7% (Fig. 5A, 6B). The observed 2.5-fold CNR increase for resting-state (from 0.13 ± 0.03 to 0.34 ± 0.6; Fig. 5B) is close to the previously reported 3-fold CNR increase achieved using a block task-paradigm (Leite et al., 2002; Vanduffel et al., 2001). However, relative to anesthetized BOLD (0.34 ± 0.06 vs. 0.21 ± 0.07; Fig. 6C), the improvement is more modest.

Despite the improvement in CNR and the rigid head-coil attachment, functional timeseries in awake macaques contain a large amount of spatially time-varying artifacts (e.g., subtle motion, and respiration) that must be removed from the data to obtain neurobiologically sensible FC profiles (Fig. 6, Fig. 9B) (Autio et al., 2021; Griffanti et al., 2014; Power et al., 2012). Indeed, motion and structured artifacts accounted for ≈24% of the variance in awake macaque CBVw resting-state fMRI, which together represent about three-fold more variance than neural signals (Fig. 6B). Comparatively, in humans, these artifacts are even more pronounced, contributing approximately seven times more variance than neural signals (Marcus et al., 2013). We used sICA-FIX to remove these motion and structured noise artifacts from the data thereby improving seed-based FC estimations (Griffanti et al., 2014). For instance, denoising had a profound effect on CBVw resting-state FC between MT and FEF (Fig. 7) (Markov et al., 2014). These results are in line with our preliminary report that have demonstrated that FIX-ICA denoising improves the comparison between quantitative retrograde tracer and FC in anesthetized macaques (Van Essen et al., 2019; Hayashi et al., 2021; Markov et al., 2014).

One known limitation of the sICA-FIX clean-up, however, is that it does not effectively remove global artifacts (Glasser et al., 2018). This is a potential concern in CBVw fMRI, where the relative global gray matter timeseries variance was notably high before ICA-FIX cleanup (≈7.4%). This global variance may reflect variations in global neural activity (Schölvinck et al., 2010), respiration and washout of the ferumoxytol contrast agent. In contrast, anesthetized macaque BOLD fMRI experiments, where respiration is externally controlled by a ventilation machine (at ≈0.2 Hz), the global variance is merely 0.2% (Autio et al., 2020, 2021). This raises questions whether anesthesia has a strong suppressive effect on global neural activity or merely reduces variability in respiration. To disentangle these physiological processes in the frequency domain, the temporal sampling rate (e.g., TR) should be increased to above the critical sampling frequency (> 2 × frequency of physiological noise). Unfortunately, temporal resolution used herein (TR = 0.7 s = 1.4 Hz) is insufficient to properly sample the cardiac cycle in awake macaques. Consequently, physiological noise from respiration (≈0.2 Hz), cardiac cycles (≈1-2 Hz), and scanner-related helium-pump artifacts (Kiviniemi et al., 2016) may be aliased into and overlap with slower gradient heating-based signal drift, vascular tone and neuronal processes (Uchiyama et al., 2007) (mainly <0.1 Hz) in the frequency domain. Harmonizing temporal resolution across species could benefit from adjusting the sampling rates according to the animal’s physiology, albeit this is impractical in smaller species with progressively faster respiration (e.g. ≈0.2, 0.2, 1.3 and 1.8 Hz in human, macaque, marmoset and rat, respectively) and cardiac cycles (≈1, 2, 5 and 7 Hz, respectively) (Baran et al., 2020; Bishop et al., 2022). To address these challenges, our approach is to acquire a large number of fMRI volumes and to selectively remove the remaining respiration- and ferumoxytol washout-related global artifacts from global neural activity at the group-level using temporal ICA (Glasser et al., 2018).

Another important consideration is the variability of FC across subjects within each species (Xu et al., 2019), which might reflect species-specific behavioral traits (Finn et al., 2015; Yokoyama et al., 2021). However, such investigations necessitate careful estimation of measurement biases. Here, we demonstrated that within-scan, within-subject, and between-subjects FC reproducibility are similar between macaques and humans (Koike et al., 2021; Laumann et al., 2015; Murphy et al., 2007). Although the awake macaque population size used here is small (N=3), these conclusions are supported by results from anesthetized macaques (N=5; (Autio et al., 2021)).

The similar inter-subject variability in FC between species was unexpected, particularly given that humans exhibit much greater variability in gyrification patterns compared to macaques (Hayashi et al., 2021). This finding suggests that neuroanatomical features like cortical folding may not be the primary drivers of variability in FC. Instead, it could suggest that neuronal activity patterns underlying FC are conserved across these species or are influenced by factors other than surface anatomy, such as macrovasculature (Autio et al., 2024; Kurzawski et al., 2022), CNR, synaptic architecture, or the intrinsic dynamics of neural circuits (Chaudhuri et al., 2015; Douglas & Martin, 2007). Indeed, neuronal high-frequency rhythms (Buzsáki et al., 2013) and neurovascular coupling mechanisms (Logothetis et al., 2001) are evolutionarily preserved and may translate into the similar temporal receptive fields and low-frequency vascular dynamics across primates.

The excellent reproducibility of the macaque data enables high-precision mapping of dense functional connectivity across sessions and subjects (Fig. 8B, 9B). In both species, we observed FC extending across sensorimotor and premotor areas (macaque: F1, 3a/b, 1, 2, 5, F2, and F5; human: M1, 3a/b, 1, 2, 7C, 6d, and 6v), the central proportion of the insula, and parietal cortex (macaque: proportions of areas SII, 7OP, and 7B; human PO). These results suggest that the dorsal proportion of macaque F5 may be a homologue of human area 6v, F2 may correspond to 6d and 5 to area 7C. In macaques, we identified two additional positively correlated nodes (9/46v and 9/46d) that were absent in the human resting-state FC maps (Fig. 6D). Anatomical and physiological evidence in macaques has linked areas 9/46v and 9/46d to complex motor planning, including arm movements (Barbas and Pandya 1989; Hoshi & Tanji, 2004), and with similar functional evidence reported in humans (Beurze et al., 2007). It remains unclear whether this discrepancy stems from differences in data acquisition methods (CBVw vs. BOLD fMRI), differences in video presentation (macaques: dynamic Inscape; humans: static crosshair), analytic limitations (e.g., multivariate analysis, surface registration), or evolutionary differences. Despite these uncertainties, these homologous FC patterns provide a strong foundation for comparative neuroimaging and improved cortical landmarks for inter-species registration.

While our primary focus has been to demonstrate the ability to measure neural activity in the cerebral cortex, fMRI also offers a powerful tool for exploring subcortical circuits (Fig. 10B, C). Notably, subcortical structures exhibit superior FC in awake macaques compared to anesthetized ones, likely due in part to the thalamus, which is particularly sensitive to anesthesia (Mapelli et al., 2021). This methodology, therefore, enhances our ability to investigate polysynaptic cortico-subcortical-cortical circuits, such as those involving the basal ganglia and cerebellar cortices—structures whose evolution remains poorly understood due to the lack of robust anatomical landmarks and the inherent challenges of studying polysynaptic circuits (Kelly & Strick, 2000; Magielse et al., 2023).

## 5. Conclusions

A 24-channel 3T receive coil was developed for accelerated neuroimaging of awake macaque monkeys. The use of accelerated ferumoxytol-weighted fMRI, combined with ICA-FIX artifact removal, produced highly reproducible parcellated functional connectivity patterns at the single-subject level. At the group-level, dense seed-based functional connectivity exhibited neurobiologically robust and likely homologous networks in macaques and humans. The data acquisition hardware, imaging protocols, and data analysis pipelines presented here are publicly available, facilitating further exploration of the functional organization of primate brains by the broader scientific community.

## Data and Code Availability

The 24-channel RF receive coil for awake macaque monkeys is commercially available via Rogue Research, Montreal, Canada https://www.rogue-research.com/; produced by Takashima Seisakusho Co. Ltd. Tokyo, Japan.

The imaging protocols are available at https://brainminds-beyond.riken.jp/hcp-nhp-protocol.

The NHP-HCP pipelines will be made available at https://github.com/Washington-University/HCPpipelines.

The RestingStateStats script is available at https://github.com/Washington-University/HCPpipelines.

## CRediT authorship contribution statement

**Joonas Autio:** Conceptualization, Methodology, Formal analysis, Investigation, Writing - original draft, Writing - review & editing, Visualization. **Atsushi Yoshida:** Formal analysis, Investigation, Visualization. **Yoshihiko Kawabata:** Methodology, Investigation. **Masahiro Ohno:** Investigation, Visualization. **Kantaro Nishigori:** Investigation. **Takayuki Ose:** Investigation, Data Curation. **Stephen Smith:** Software, Writing - review & editing. **David Van Essen:** Writing - review & editing, Funding acquisition. **Matthew Glasser:** Conceptualization, Software, Formal analysis, Funding acquisition. **Takuya Hayashi:** Conceptualization, Methodology, Software, Formal analysis, Writing - review & editing, Resources, Funding acquisition.

## Acknowledgements

The authors appreciate technical contributions and discussions from Takashi Azuma, Erin Reid, Paul McCarthy, Katsutos’hi Murata, Yuta Urushibata, Takuro Ikeda, Masataka Yamaguchi, Yuki Hori, Akihiro Kawasaki, Chihiro Takeda and Reiko Kobayashi.

## 6. Appendices

### 6.1. Data Acquisition Strategy

To improve comparison of brain organization across primates, our data acquisition strategy is based on standardized minimal requirements, adjusting image resolution to the thinnest parts of the cerebral cortex (Autio et al., 2020; Glasser et al., 2013; Hayashi et al., 2021; Hori et al., 2018). The structural image resolution (0.5 mm) is approximately scaled to half of the minimum cortical thickness in macaques (≈1 mm) to ensure accurate reconstruction of the cortical pial and white matter surfaces. The fMRI spatial resolution (1.25 mm) is scaled below the 5^th^ percentile of cortical thickness (≈1.4 mm) to reduce partial volume effects from cerebrospinal fluid and white matter, and to improve differentiation between opposing banks of sulci (Autio et al., 2020; Glasser et al., 2013).

While our previous BOLD resting-state fMRI image acquisition protocol in anesthetized macaques closely followed YA-HCP’s acquisition protocols at 3T (Autio et al., 2020; Glasser et al., 2013), CBVw fMRI data acquisition in awake macaques requires different methodological considerations (Autio et al., 2021; Leite et al., 2002; Vanduffel et al., 2001). First, scan durations are more limited in awake macaques, necessitating the acquisition of data with high temporal sampling rates (e.g. short TR) for robust detection of functional networks. Second, in CBVw fMRI shorter TEs matched with contrast agent dose induced T_2_*=TE enhances contrast (see below) (Mandeville et al., 2004).

The intravascular injection of magnetic susceptibility contrast agent (e.g. ferumoxytol) effectively attenuates the intravascular blood water compartment and CSF nuisance signals located adjacent to major arteries and draining veins along the cortical surface (Autio, et al., 2024; Zhao et al., 2006). Thus, the gray matter transverse relaxation rate (R_2_=1/T_2_) in CBVw fMRI can be expressed as a single-compartment model:

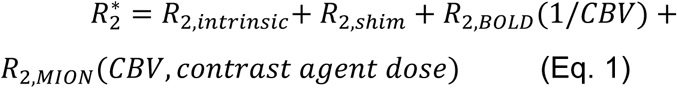

where R_2,intrinsic_ corresponds to intrinsic relaxation processes, R_2,shim_ represents susceptibility-induced B_0_ field inhomogeneities (e.g., near air-tissue interfaces), and R_2,BOLD_ is (extravascular) blood oxygen level dependent (BOLD) contrast, which is inversely proportional to CBV. Conversely, R_2,ferumoxytol_ is directly proportional to CBV (Mandeville et al., 2004).

In Eq. 1, signal loss due to the first three relaxation terms is of no interest and should be minimized, while the last contrast agent dose-dependent term should be maximized. Practically, this is achieved by minimizing TE and then optimizing the contrast agent dosage, thereby enhancing the relative contribution of R_2,ferumoxytol_ (Mandeville et al., 2004). At spatial resolutions (1.25 mm isotropic), where the venous and arterial vessel contributions are not separable (Autio, et al., 2024; Weber et al., 2008), it is crucial to use a contrast agent dose that overcomes the BOLD effect (so that R_2,ferumoxytol_ >> R_2,BOLD_), while maintaining sufficient tSNR to detect neural activity-associated signal fluctuations. For simplicity, assume the noise (σ_N_) is constant across image, then the dynamic resting-state fMRI CNR can be expressed as:

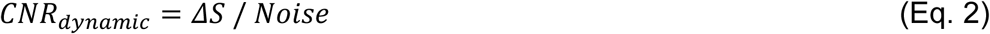

Given the ΔR_2,ferumoxytol_ is small (due to the < 10% CBV changes), one can approximate the exponential difference as:

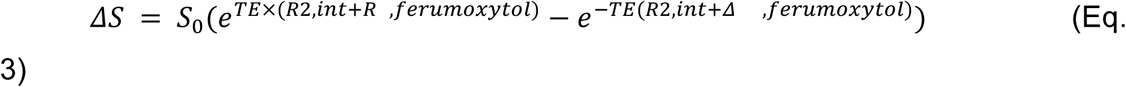

Using a first-order Taylor expansion for small ΔR_2,ferumoxytol_:

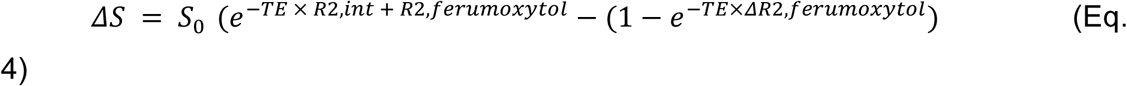

Taking the derivative with respect to CBV and setting it to zero:

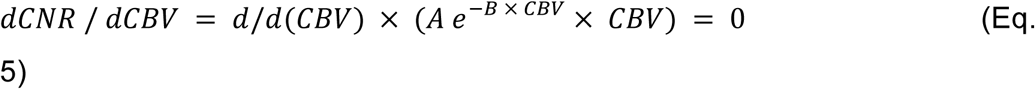

We acquire optimal contrast agent dose:

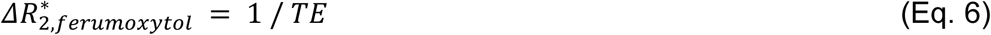

This theoretical calculation, based on Bloch equations and a single-tissue model, predicts optimal CNR using intravascular dose of ferumoxytol that induces change in 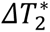 that matches the (minimal) TE.

**Supplementary Figure 1.**
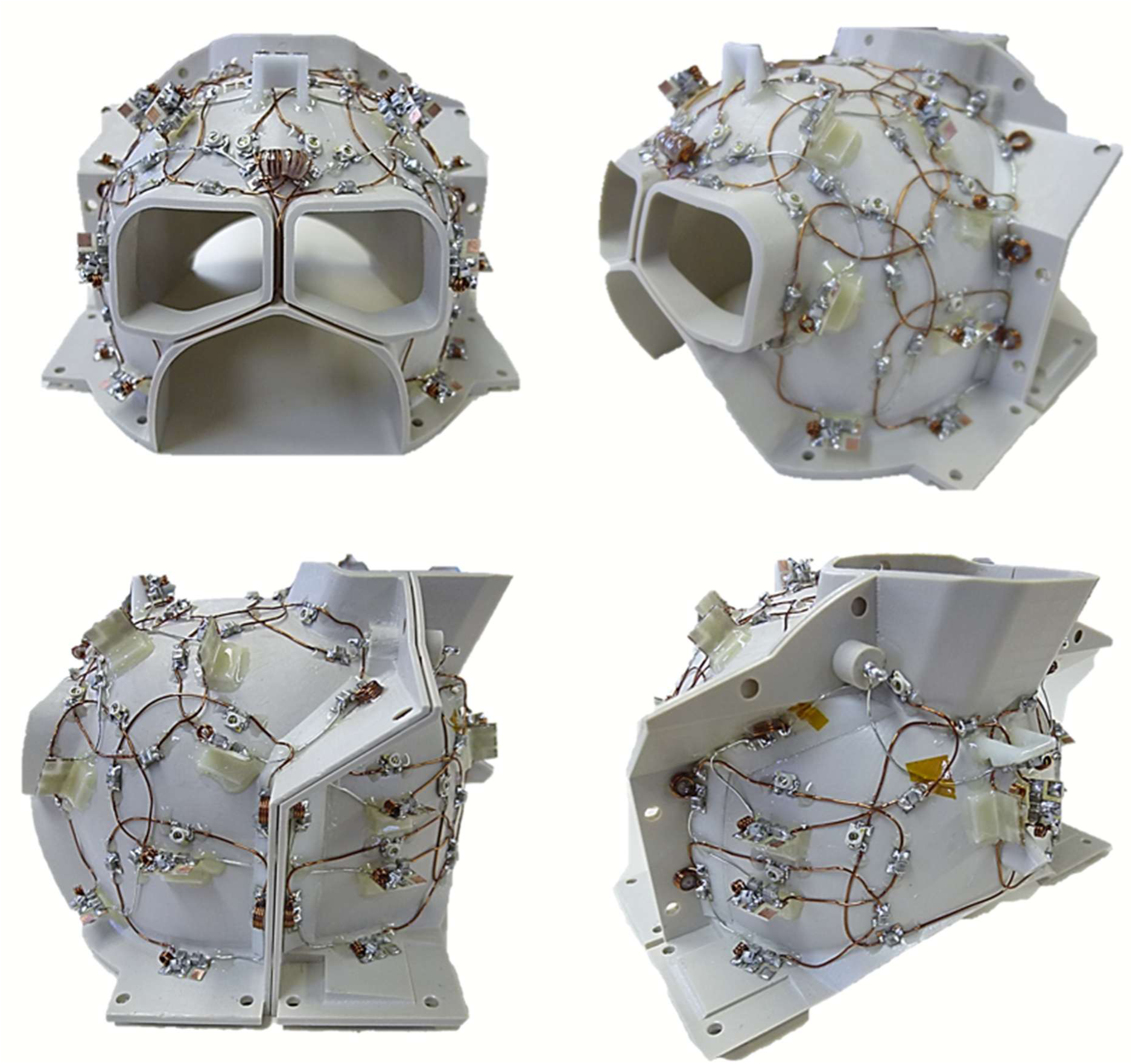
Additional views of the 24-channel phased-array macaque coil.

**Supplementary Figure 2.**
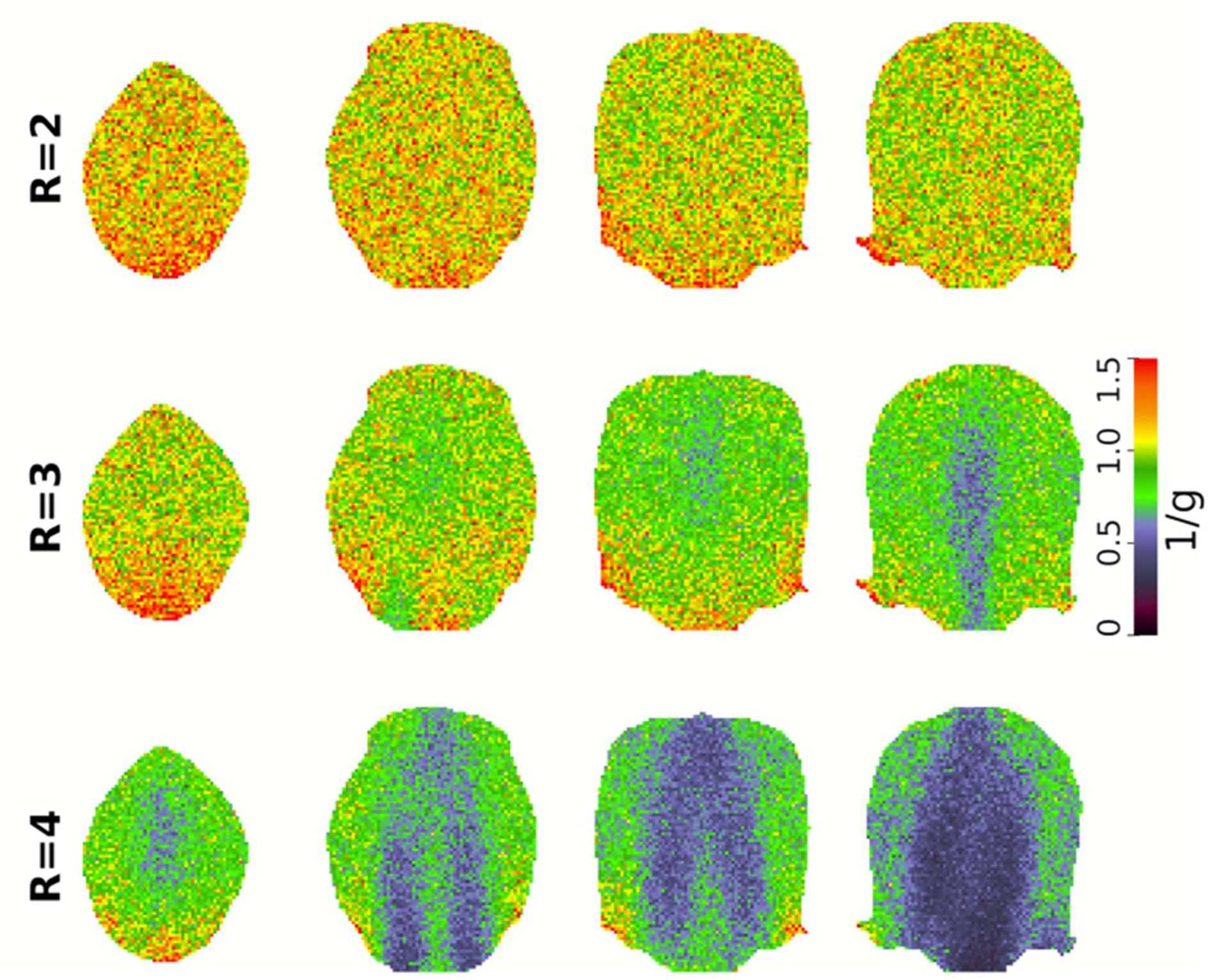
Geometry (g)-factor quality assessment for different acceleration factors. Reduction factors (R) of 2 and 3 show reasonable geometry-dependent signal losses, whereas R = 4 results in lower 1/g values, implying greater noise amplification during image reconstruction.

**Supplementary Figure 3.**
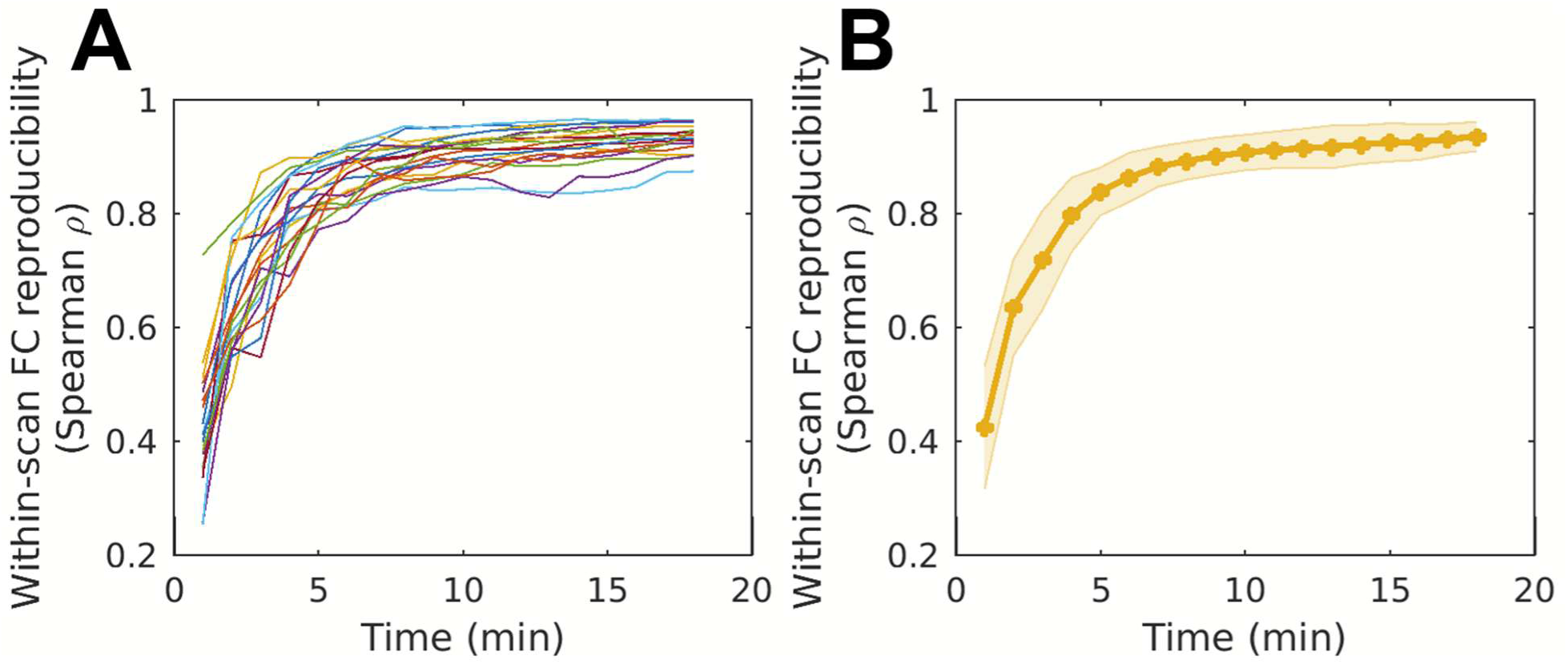
Within-scan reproducibility of M132 parcellated functional connectivity matrix. **(A)** Reproducibility of within-scan M132 atlas-parcellated functional connectivity correlation matrix plotted as a function of scan duration. Each color indicates a separate scan session. **(B)** On average, within scan reproducibility reached excellent quality (Spearman Rho > 0.90) using a pair of 10 minute scans.

**Supplementary Figure 4.**
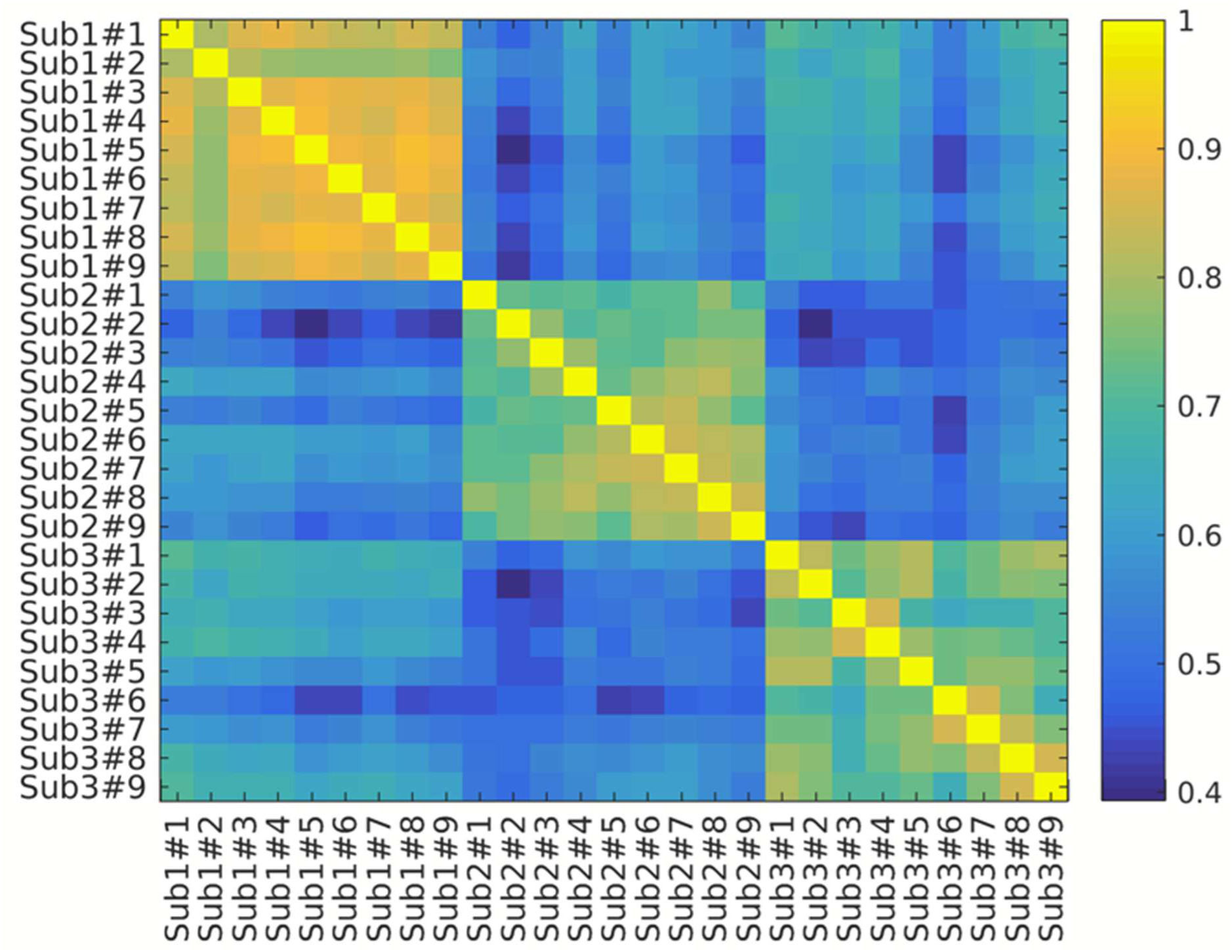
Identifying subjects using noise-dominant correlation patterns.

## References

Andersson, J. L. R., Skare, S., & Ashburner, J. (2003). How to correct susceptibility distortions in spin-echo echo-planar images: Application to diffusion tensor imaging. NeuroImage, 20(2), Article 2. 10.1016/S1053-8119(03)00336-7

Autio, J. A., Glasser, M. F., Ose, T., Donahue, C. J., Bastiani, M., Ohno, M., Kawabata, Y., Urushibata, Y., Murata, K., Nishigori, K., Yamaguchi, M., Hori, Y., Yoshida, A., Go, Y., Coalson, T. S., Jbabdi, S., Sotiropoulos, S. N., Kennedy, H., Smith, S., … Hayashi, T. (2020). Towards HCP-Style macaque connectomes: 24-Channel 3T multi-array coil, MRI sequences and preprocessing. NeuroImage, 215, 116800. 10.1016/j.neuroimage.2020.116800

Autio, J. A., Kimura, I., Ose, T., Matsumoto, Y., Ohno, M., Urushibata, Y., Ikeda, T., Glasser, M. F., Essen, D. C. V., & Hayashi, T. (2024). Mapping vascular network architecture in primate brain using ferumoxytol-weighted laminar MRI. eLife, 13. 10.7554/eLife.99940.1

Autio, J. A., Uematsu, A., Ikeda, T., Ose, T., Hou, Y., Magrou, L., Kimura, I., Ohno, M., Murata, K., Coalson, T., Kennedy, H., Glasser, M. F., Essen, D. C. V., & Hayashi, T. (2024). Charting Cortical-Layer Specific Area Boundaries Using Gibbs Ringing Attenuated T1w/T2w-FLAIR Myelin MRI (p. 2024.09.27.615294). bioRxiv. 10.1101/2024.09.27.615294

Autio, J. A., Zhu, Q., Li, X., Glasser, M. F., Schwiedrzik, C. M., Fair, D. A., Zimmermann, J., Yacoub, E., Menon, R. S., Van Essen, D. C., Hayashi, T., Russ, B., & Vanduffel, W. (2021). Minimal specifications for non-human primate MRI: Challenges in standardizing and harmonizing data collection. NeuroImage, 236, 118082. 10.1016/j.neuroimage.2021.118082

Baran, S. W., Gupta, A. D., Lim, M. A., Mathur, A., Rowlands, D. J., Schaevitz, L. R., Shanmukhappa, S. K., & Walker, D. B. (2020). Continuous, Automated Breathing Rate and Body Motion Monitoring of Rats With Paraquat-Induced Progressive Lung Injury. Frontiers in Physiology, 11, 569001. 10.3389/fphys.2020.569001

Birn, R. M., Molloy, E. K., Patriat, R., Parker, T., Meier, T. B., Kirk, G. R., Nair, V. A., Meyerand, M. E., & Prabhakaran, V. (2013). The effect of scan length on the reliability of resting-state fMRI connectivity estimates. NeuroImage, 83, 550–558. 10.1016/j.neuroimage.2013.05.099

Bishop, M., Weinhold, M., Turk, A. Z., Adeck, A., & SheikhBahaei, S. (2022). An open-source tool for automated analysis of breathing behaviors in common marmosets and rodents. eLife, 11, e71647. 10.7554/eLife.71647

Botvinik-Nezer, R., Holzmeister, F., Camerer, C. F., Dreber, A., Huber, J., Johannesson, M., Kirchler, M., Iwanir, R., Mumford, J. A., Adcock, R. A., Avesani, P., Baczkowski, B. M., Bajracharya, A., Bakst, L., Ball, S., Barilari, M., Bault, N., Beaton, D., Beitner, J., … Schonberg, T. (2020). Variability in the analysis of a single neuroimaging dataset by many teams. Nature, 582(7810), Article 7810. 10.1038/s41586-020-2314-9

Boxerman, J. L., Hamberg, L. M., Rosen, B. R., & Weisskoff, R. M. (1995). Mr contrast due to intravascular magnetic susceptibility perturbations. Magnetic Resonance in Medicine, 34(4), Article 4. 10.1002/mrm.1910340412

Buzsáki, G., Logothetis, N., & Singer, W. (2013). Scaling Brain Size, Keeping Timing: Evolutionary Preservation of Brain Rhythms. Neuron, 80(3), Article 3. 10.1016/j.neuron.2013.10.002

Chaudhuri, R., Knoblauch, K., Gariel, M.-A., Kennedy, H., & Wang, X.-J. (2015). A large-scale circuit mechanism for hierarchical dynamical processing in the primate cortex. Neuron, 88(2), Article 2. 10.1016/j.neuron.2015.09.008

Chung, S., Kim, D., Breton, E., & Axel, L. (2010). Rapid B1+ mapping using a preconditioning RF pulse with TurboFLASH readout. Magnetic Resonance in Medicine, 64(2), Article 2. 10.1002/mrm.22423

Donahue, C. J., Glasser, M. F., Preuss, T. M., Rilling, J. K., & Essen, D. C. V. (2018). Quantitative assessment of prefrontal cortex in humans relative to nonhuman primates. Proceedings of the National Academy of Sciences, 201721653. 10.1073/pnas.1721653115

Douglas, R. J., & Martin, K. A. C. (2007). Recurrent neuronal circuits in the neocortex. Current Biology, 17(13), Article 13. 10.1016/j.cub.2007.04.024

Finn, E. S., Shen, X., Scheinost, D., Rosenberg, M. D., Huang, J., Chun, M. M., Papademetris, X., & Constable, R. T. (2015). Functional connectome fingerprinting: Identifying individuals using patterns of brain connectivity. Nature Neuroscience, 18(11), Article 11. 10.1038/nn.4135

Fischl, B. (2012). FreeSurfer. NeuroImage, 62(2), 774–781. 10.1016/j.neuroimage.2012.01.021

Gilbert, K. M., Gati, J. S., Barker, K., Everling, S., & Menon, R. S. (2016). Optimized parallel transmit and receive radiofrequency coil for ultrahigh-field MRI of monkeys. NeuroImage, 125, 153–161. 10.1016/j.neuroimage.2015.10.048

Glasser, M. F., Coalson, T. S., Bijsterbosch, J. D., Harrison, S. J., Harms, M. P., Anticevic, A., Van Essen, D. C., & Smith, S. M. (2018). Using temporal ICA to selectively remove global noise while preserving global signal in functional MRI data. NeuroImage, 181, 692–717. 10.1016/j.neuroimage.2018.04.076

Glasser, M. F., Coalson, T. S., Robinson, E. C., Hacker, C. D., Harwell, J., Yacoub, E., Ugurbil, K., Andersson, J., Beckmann, C. F., Jenkinson, M., Smith, S. M., & Van Essen, D. C. (2016). A multi-modal parcellation of human cerebral cortex. Nature, 536(7615), Article 7615. 10.1038/nature18933

Glasser, M. F., Sotiropoulos, S. N., Wilson, J. A., Coalson, T. S., Fischl, B., Andersson, J. L., Xu, J., Jbabdi, S., Webster, M., Polimeni, J. R., Van Essen, D. C., & Jenkinson, M. (2013). The minimal preprocessing pipelines for the Human Connectome Project. NeuroImage, 80, 105–124. 10.1016/j.neuroimage.2013.04.127

Greve, D. N., & Fischl, B. (2009). Accurate and robust brain image alignment using boundary-based registration. NeuroImage, 48(1), Article 1. 10.1016/j.neuroimage.2009.06.060

Griffanti, L., Douaud, G., Bijsterbosch, J., Evangelisti, S., Alfaro-Almagro, F., Glasser, M. F., Duff, E. P., Fitzgibbon, S., Westphal, R., Carone, D., Beckmann, C. F., & Smith, S. M. (2017). Hand classification of fMRI ICA noise components. Neuroimage, 154, 188–205. 10.1016/j.neuroimage.2016.12.036

Griffanti, L., Salimi-Khorshidi, G., Beckmann, C. F., Auerbach, E. J., Douaud, G., Sexton, C. E., Zsoldos, E., Ebmeier, K. P., Filippini, N., Mackay, C. E., Moeller, S., Xu, J., Yacoub, E., Baselli, G., Ugurbil, K., Miller, K. L., & Smith, S. M. (2014). ICA-based artefact removal and accelerated fMRI acquisition for improved resting state network imaging. NeuroImage, 95, 232–247. 10.1016/j.neuroimage.2014.03.034

Griswold, M. A., Jakob, P. M., Heidemann, R. M., Nittka, M., Jellus, V., Wang, J., Kiefer, B., & Haase, A. (2002). Generalized autocalibrating partially parallel acquisitions (GRAPPA). Magnetic Resonance in Medicine, 47(6), Article 6. 10.1002/mrm.10171

Hayashi, T., Hou, Y., Glasser, M. F., Autio, J. A., Knoblauch, K., Inoue-Murayama, M., Coalson, T., Yacoub, E., Smith, S., Kennedy, H., & Van Essen, D. C. (2021). The nonhuman primate neuroimaging and neuroanatomy project. NeuroImage, 229, 117726. 10.1016/j.neuroimage.2021.117726

Hori, Y., Autio, J. A., Ohno, M., Kawabata, Y., Urushibata, Y., Murata, K., Yamaguchi, M., Kawasaki, A., Takeda, C., Yokoyama, C., Glasser, M. F., & Hayashi, T. (2018). Translating the Human Connectome Project to Marmoset Imaging: 16-Channel Multi-Array Coil and HCP-Style MRI Protocols and Preprocessing. https://archive.ismrm.org/2018/3404.html

Hutchison, R. M., & Everling, S. (2012). Monkey in the middle: Why non-human primates are needed to bridge the gap in resting-state investigations. Frontiers in Neuroanatomy, 6. 10.3389/fnana.2012.00029

Hutchison, R. M., Leung, L. S., Mirsattari, S. M., Gati, J. S., Menon, R. S., & Everling, S. (2011). Resting-state networks in the macaque at 7 T. NeuroImage, 56(3), Article 3. 10.1016/j.neuroimage.2011.02.063

Jenkinson, M., Bannister, P., Brady, M., & Smith, S. (2002). Improved Optimization for the Robust and Accurate Linear Registration and Motion Correction of Brain Images. NeuroImage, 17(2), Article 2. 10.1006/nimg.2002.1132

Jenkinson, M., Beckmann, C. F., Behrens, T. E. J., Woolrich, M. W., & Smith, S. M. (2012). FSL. NeuroImage, 62(2), Article 2. 10.1016/j.neuroimage.2011.09.015

Kelly, R. M., & Strick, P. L. (2000). Rabies as a transneuronal tracer of circuits in the central nervous system. Journal of Neuroscience Methods, 103(1), 63–71. 10.1016/s0165-0270(00)00296-x

Kiviniemi, V., Wang, X., Korhonen, V., Keinänen, T., Tuovinen, T., Autio, J., LeVan, P., Keilholz, S., Zang, Y.-F., Hennig, J., & Nedergaard, M. (2016). Ultra-fast magnetic resonance encephalography of physiological brain activity – Glymphatic pulsation mechanisms? Journal of Cerebral Blood Flow & Metabolism, 36(6), Article 6. 10.1177/0271678X15622047

Koike, S., Tanaka, S. C., Okada, T., Aso, T., Yamashita, A., Yamashita, O., Asano, M., Maikusa, N., Morita, K., Okada, N., Fukunaga, M., Uematsu, A., Togo, H., Miyazaki, A., Murata, K., Urushibata, Y., Autio, J., Ose, T., Yoshimoto, J., … Hayashi, T. (2021). Brain/MINDS beyond human brain MRI project: A protocol for multi-level harmonization across brain disorders throughout the lifespan. NeuroImage: Clinical, 102600. 10.1016/j.nicl.2021.102600

Kurzawski, J. W., Gulban, O. F., Jamison, K., Winawer, J., & Kay, K. (2022). Non-Neural Factors Influencing BOLD Response Magnitudes within Individual Subjects. Journal of Neuroscience, 42(38), 7256–7266. 10.1523/JNEUROSCI.2532-21.2022

Laumann, T. O., Gordon, E. M., Adeyemo, B., Snyder, A. Z., Joo, S. J., Chen, M.-Y., Gilmore, A. W., McDermott, K. B., Nelson, S. M., Dosenbach, N. U. F., Schlaggar, B. L., Mumford, J. A., Poldrack, R. A., & Petersen, S. E. (2015). Functional System and Areal Organization of a Highly Sampled Individual Human Brain. Neuron, 87(3), Article 3. 10.1016/j.neuron.2015.06.037

Leite, F. P., Tsao, D., Vanduffel, W., Fize, D., Sasaki, Y., Wald, L. L., Dale, A. M., Kwong, K. K., Orban, G. A., Rosen, B. R., Tootell, R. B. H., & Mandeville, J. B. (2002). Repeated fMRI Using Iron Oxide Contrast Agent in Awake, Behaving Macaques at 3 Tesla. NeuroImage, 16(2), Article 2. 10.1006/nimg.2002.1110

Logothetis, N. K., Pauls, J., Augath, M., Trinath, T., & Oeltermann, A. (2001). Neurophysiological investigation of the basis of the fMRI signal. Nature, 412(6843), 150–157. 10.1038/35084005

Magielse, N., Toro, R., Steigauf, V., Abbaspour, M., Eickhoff, S. B., Heuer, K., & Valk, S. L. (2023). Phylogenetic comparative analysis of the cerebello-cerebral system in 34 species highlights primate-general expansion of cerebellar crura I-II. Communications Biology, 6(1), 1–17. 10.1038/s42003-023-05553-z

Mandeville, J. B., Jenkins, B. G., Chen, Y.-C. I., Choi, J.-K., Kim, Y. R., Belen, D., Liu, C., Kosofsky, B. E., & Marota, J. J. A. (2004). Exogenous contrast agent improves sensitivity of gradient-echo functional magnetic resonance imaging at 9.4 T. Magnetic Resonance in Medicine, 52(6), Article 6. 10.1002/mrm.20278

Mandeville, J. B., Marota, J. J. A., Ayata, C., Moskowitz, M. A., Weisskoff, R. M., & Rosen, B. R. (1999). MRI measurement of the temporal evolution of relative CMRO2 during rat forepaw stimulation. Magnetic Resonance in Medicine, 42(5), 944–951. 10.1002/(SICI)1522-2594(199911)42:5<944::AID-MRM15>3.0.CO;2-W

Mantini, D., Gerits, A., Nelissen, K., Durand, J.-B., Joly, O., Simone, L., Sawamura, H., Wardak, C., Orban, G. A., Buckner, R. L., & Vanduffel, W. (2011). Default mode of brain function in monkeys. The Journal of Neuroscience: The Official Journal of the Society for Neuroscience, 31(36), Article 36. 10.1523/JNEUROSCI.2318-11.2011

Mapelli, J., Gandolfi, D., Giuliani, E., Casali, S., Congi, L., Barbieri, A., D’Angelo, E., & Bigiani, A. (2021). The effects of the general anesthetic sevoflurane on neurotransmission: An experimental and computational study. Scientific Reports, 11(1), 4335. 10.1038/s41598-021-83714-y

Marcus, D. S., Harms, M. P., Snyder, A. Z., Jenkinson, M., Wilson, J. A., Glasser, M. F., Barch, D. M., Archie, K. A., Burgess, G. C., Ramaratnam, M., Hodge, M., Horton, W., Herrick, R., Olsen, T., McKay, M., House, M., Hileman, M., Reid, E., Harwell, J., … Van Essen, D. C. (2013). Human Connectome Project informatics: Quality control, database services, and data visualization. NeuroImage, 80, 202–219. 10.1016/j.neuroimage.2013.05.077

Markov, N. T., Ercsey-Ravasz, M. M., Ribeiro Gomes, A. R., Lamy, C., Magrou, L., Vezoli, J., Misery, P., Falchier, A., Quilodran, R., Gariel, M. A., Sallet, J., Gamanut, R., Huissoud, C., Clavagnier, S., Giroud, P., Sappey-Marinier, D., Barone, P., Dehay, C., Toroczkai, Z., … Kennedy, H. (2014). A Weighted and Directed Interareal Connectivity Matrix for Macaque Cerebral Cortex. Cerebral Cortex, 24(1), Article 1. 10.1093/cercor/bhs270

Milham, M. P., Ai, L., Koo, B., Xu, T., Amiez, C., Balezeau, F., Baxter, M. G., Blezer, E. L. A., Brochier, T., Chen, A., Croxson, P. L., Damatac, C. G., Dehaene, S., Everling, S., Fair, D. A., Fleysher, L., Freiwald, W., Froudist-Walsh, S., Griffiths, T. D., … Schroeder, C. E. (2018). An Open Resource for Non-human Primate Imaging. Neuron, 100(1), Article 1. 10.1016/j.neuron.2018.08.039

Moeller, S., Yacoub, E., Olman, C. A., Auerbach, E., Strupp, J., Harel, N., & Uğurbil, K. (2010). Multiband multislice GE-EPI at 7 tesla, with 16-fold acceleration using partial parallel imaging with application to high spatial and temporal whole-brain fMRI. Magnetic Resonance in Medicine, 63(5), Article 5. 10.1002/mrm.22361

Murphy, K., Bodurka, J., & Bandettini, P. A. (2007). How long to scan? The relationship between fMRI temporal signal to noise ratio and necessary scan duration. NeuroImage, 34(2), Article 2. 10.1016/j.neuroimage.2006.09.032

Nickerson, L. D. (2018). Replication of Resting State-Task Network Correspondence and Novel Findings on Brain Network Activation During Task fMRI in the Human Connectome Project Study. Scientific Reports, 8(1), 17543. 10.1038/s41598-018-35209-6

Ogawa, S., Menon, R. S., Tank, D. W., Kim, S. G., Merkle, H., Ellermann, J. M., & Ugurbil, K. (1993). Functional brain mapping by blood oxygenation level-dependent contrast magnetic resonance imaging. A comparison of signal characteristics with a biophysical model. Biophysical Journal, 64(3), Article 3.

Ose, T., Autio, J. A., Ohno, M., Frey, S., Uematsu, A., Kawasaki, A., Takeda, C., Hori, Y., Nishigori, K., Nakako, T., Yokoyama, C., Nagata, H., Yamamori, T., Van Essen, D. C., Glasser, M. F., Watabe, H., & Hayashi, T. (2022). Anatomical variability, multi-modal coordinate systems, and precision targeting in the marmoset brain. NeuroImage, 250, 118965. 10.1016/j.neuroimage.2022.118965

Paasonen, J., Stenroos, P., Salo, R. A., Kiviniemi, V., & Gröhn, O. (2018). Functional connectivity under six anesthesia protocols and the awake condition in rat brain. NeuroImage, 172, 9–20. 10.1016/j.neuroimage.2018.01.014

Polimeni, J. R., Bhat, H., Witzel, T., Benner, T., Feiweier, T., Inati, S. J., Renvall, V., Heberlein, K., & Wald, L. L. (2016). Reducing sensitivity losses due to respiration and motion in accelerated echo planar imaging by reordering the autocalibration data acquisition. Magnetic Resonance in Medicine, 75(2), Article 2. 10.1002/mrm.25628

Power, J. D., Barnes, K. A., Snyder, A. Z., Schlaggar, B. L., & Petersen, S. E. (2012). Spurious but systematic correlations in functional connectivity MRI networks arise from subject motion. NeuroImage, 59(3), Article 3. 10.1016/j.neuroimage.2011.10.018

Robinson, E. C., Garcia, K., Glasser, M. F., Chen, Z., Coalson, T. S., Makropoulos, A., Bozek, J., Wright, R., Schuh, A., Webster, M., Hutter, J., Price, A., Cordero Grande, L., Hughes, E., Tusor, N., Bayly, P. V., Van Essen, D. C., Smith, S. M., Edwards, A. D., … Rueckert, D. (2018). Multimodal surface matching with higher-order smoothness constraints. NeuroImage, 167, 453–465. 10.1016/j.neuroimage.2017.10.037

Salimi-Khorshidi, G., Douaud, G., Beckmann, C. F., Glasser, M. F., Griffanti, L., & Smith, S. M. (2014). Automatic denoising of functional MRI data: Combining independent component analysis and hierarchical fusion of classifiers. NeuroImage, 90, 449–468. 10.1016/j.neuroimage.2013.11.046

Schölvinck, M. L., Maier, A., Ye, F. Q., Duyn, J. H., & Leopold, D. A. (2010). Neural basis of global resting-state fMRI activity. Proceedings of the National Academy of Sciences, 107(22), Article 22. 10.1073/pnas.0913110107

Smith, S. M., Beckmann, C. F., Andersson, J., Auerbach, E. J., Bijsterbosch, J., Douaud, G., Duff, E., Feinberg, D. A., Griffanti, L., Harms, M. P., & others. (2013). Resting-state fMRI in the human connectome project. Neuroimage, 80, 144–168.

Smith, S. M., Fox, P. T., Miller, K. L., Glahn, D. C., Fox, P. M., Mackay, C. E., Filippini, N., Watkins, K. E., Toro, R., Laird, A. R., & Beckmann, C. F. (2009). Correspondence of the brain’s functional architecture during activation and rest. Proceedings of the National Academy of Sciences, 106(31), Article 31. 10.1073/pnas.0905267106

Smith, S. M., Jenkinson, M., Woolrich, M. W., Beckmann, C. F., Behrens, T. E. J., Johansen-Berg, H., Bannister, P. R., De Luca, M., Drobnjak, I., Flitney, D. E., Niazy, R. K., Saunders, J., Vickers, J., Zhang, Y., De Stefano, N., Brady, J. M., & Matthews, P. M. (2004). Advances in functional and structural MR image analysis and implementation as FSL. NeuroImage, 23, S208–S219. 10.1016/j.neuroimage.2004.07.051

Van Dijk, K. R. A., Sabuncu, M. R., & Buckner, R. L. (2012). The influence of head motion on intrinsic functional connectivity MRI. NeuroImage, 59(1), Article 1. 10.1016/j.neuroimage.2011.07.044

van Essen, D. C., Hayashi, T., Autio, J., Ohno, M., Ose, T., Nishigori, K., Coalson, T. S., Hou, Y., Smith, S., Shen, Z., Knoblauch, K., Kennedy, H., & Glasser, M. (2019, June). Evaluation of Functional Connectivity Using Retrograde Tracers in the Macaque Monkey. OHBM. https://hal.science/hal-04968395

Vanderwal, T., Eilbott, J., & Castellanos, F. X. (2019). Movies in the magnet: Naturalistic paradigms in developmental functional neuroimaging. Developmental Cognitive Neuroscience, 36, 100600. 10.1016/j.dcn.2018.10.004

Vanderwal, T., Kelly, C., Eilbott, J., Mayes, L. C., & Castellanos, F. X. (2015). Inscapes: A movie paradigm to improve compliance in functional magnetic resonance imaging. NeuroImage, 122, 222–232. 10.1016/j.neuroimage.2015.07.069

Vanduffel, W., Fize, D., Mandeville, J. B., Nelissen, K., Van Hecke, P., Rosen, B. R., Tootell, R. B. H., & Orban, G. A. (2001). Visual Motion Processing Investigated Using Contrast Agent-Enhanced fMRI in Awake Behaving Monkeys. Neuron, 32(4), Article 4. 10.1016/S0896-6273(01)00502-5

Vincent, J. L., Patel, G. H., Fox, M. D., Snyder, A. Z., Baker, J. T., Van Essen, D. C., Zempel, J. M., Snyder, L. H., Corbetta, M., & Raichle, M. E. (2007). Intrinsic functional architecture in the anaesthetized monkey brain. Nature, 447(7140), Article 7140. 10.1038/nature05758

Vu, A. T., Jamison, K., Glasser, M. F., Smith, S. M., Coalson, T., Moeller, S., Auerbach, E. J., Ugurbil, K., & Yacoub, E. (2017). Tradeoffs in pushing the spatial resolution of fMRI for the 7 T Human Connectome Project. NeuroImage, 154, 23–32. 10.1016/j.neuroimage.2016.11.049

Weber, B., Keller, A. L., Reichold, J., & Logothetis, N. K. (2008). The Microvascular System of the Striate and Extrastriate Visual Cortex of the Macaque. Cerebral Cortex, 18(10), Article 10. 10.1093/cercor/bhm259

Wiggins, G. C., Polimeni, J. R., Potthast, A., Schmitt, M., Alagappan, V., & Wald, L. L. (2009). 96-Channel receive-only head coil for 3 Tesla: Design optimization and evaluation. Magnetic Resonance in Medicine: Official Journal of the Society of Magnetic Resonance in Medicine / Society of Magnetic Resonance in Medicine, 62(3), Article 3. 10.1002/mrm.22028

Wiggins, G. c., Triantafyllou, C., Potthast, A., Reykowski, A., Nittka, M., & Wald, L. l. (2006). 32-channel 3 Tesla receive-only phased-array head coil with soccer-ball element geometry. Magnetic Resonance in Medicine, 56(1), Article 1. 10.1002/mrm.20925

Xu, T., Nenning, K.-H., Schwartz, E., Hong, S.-J., Vogelstein, J. T., Goulas, A., Fair, D. A., Schroeder, C. E., Margulies, D. S., Smallwood, J., Milham, M. P., & Langs, G. (2020). Cross-species functional alignment reveals evolutionary hierarchy within the connectome. NeuroImage, 223, 117346. 10.1016/j.neuroimage.2020.117346

Xu, T., Sturgeon, D., Ramirez, J. S. B., Froudist-Walsh, S., Margulies, D. S., Schroeder, C. E., Fair, D. A., & Milham, M. P. (2019). Interindividual Variability of Functional Connectivity in Awake and Anesthetized Rhesus Macaque Monkeys. Biological Psychiatry: Cognitive Neuroscience and Neuroimaging, 4(6), 543–553. 10.1016/j.bpsc.2019.02.005

Yokoyama, C., Autio, J. A., Ikeda, T., Sallet, J., Mars, R. B., Van Essen, D. C., Glasser, M. F., Sadato, N., & Hayashi, T. (2021). Comparative connectomics of the primate social brain. NeuroImage, 245, 118693. 10.1016/j.neuroimage.2021.118693

Zhang, Y., Brady, M., & Smith, S. (2001). Segmentation of brain MR images through a hidden Markov random field model and the expectation-maximization algorithm. IEEE Transactions on Medical Imaging, 20(1), Article 1. 10.1109/42.906424

Zhao, F., Wang, P., Hendrich, K., Ugurbil, K., & Kim, S.-G. (2006). Cortical layer-dependent BOLD and CBV responses measured by spin-echo and gradient-echo fMRI: Insights into hemodynamic regulation. NeuroImage, 30(4), Article 4. 10.1016/j.neuroimage.2005.11.013

